# Individual variations of the human corticospinal tract and its hand-related motor fibers using diffusion MRI tractography

**DOI:** 10.1101/369199

**Authors:** K. Dalamagkas, M. Tsintou, Y. Rathi, L.J. O’Donnell, O. Pasternak, X. Gong, A. Zhu, P. Savadjiev, G.M. Papadimitriou, M. Kubicki, E.H. Yeterian, N. Makris

## Abstract

The corticospinal tract (CST) is one of the most well-studied tracts in human neuroanatomy. Its clinical significance can be demonstrated in many notable traumatic conditions and diseases such as stroke, spinal cord injury (SCI) or amyotrophic lateral sclerosis (ALS). With the advent of diffusion MRI and tractography the computational representation of the human CST in a 3D model became available. However, the representation of the entire CST and, specifically, the hand motor area has remained elusive. In this paper we proposed a novel method, using manually-drawn ROIs based on robustly identifiable neuroanatomic structures to delineate the entire CST and isolate its hand motor representation as well as to estimate their variability and generate a database of their volume, length and biophysical parameters. Using 37 healthy human subjects we performed a qualitative and quantitative analysis of the CST and the hand-related motor fiber tracts (HMFTs). Finally, we have created variability heatmaps from 37 subjects for both the aforementioned tracts, which could be utilized as reference for clinicians to explore neuropathology in both trauma and disease states.

Graphical abstract

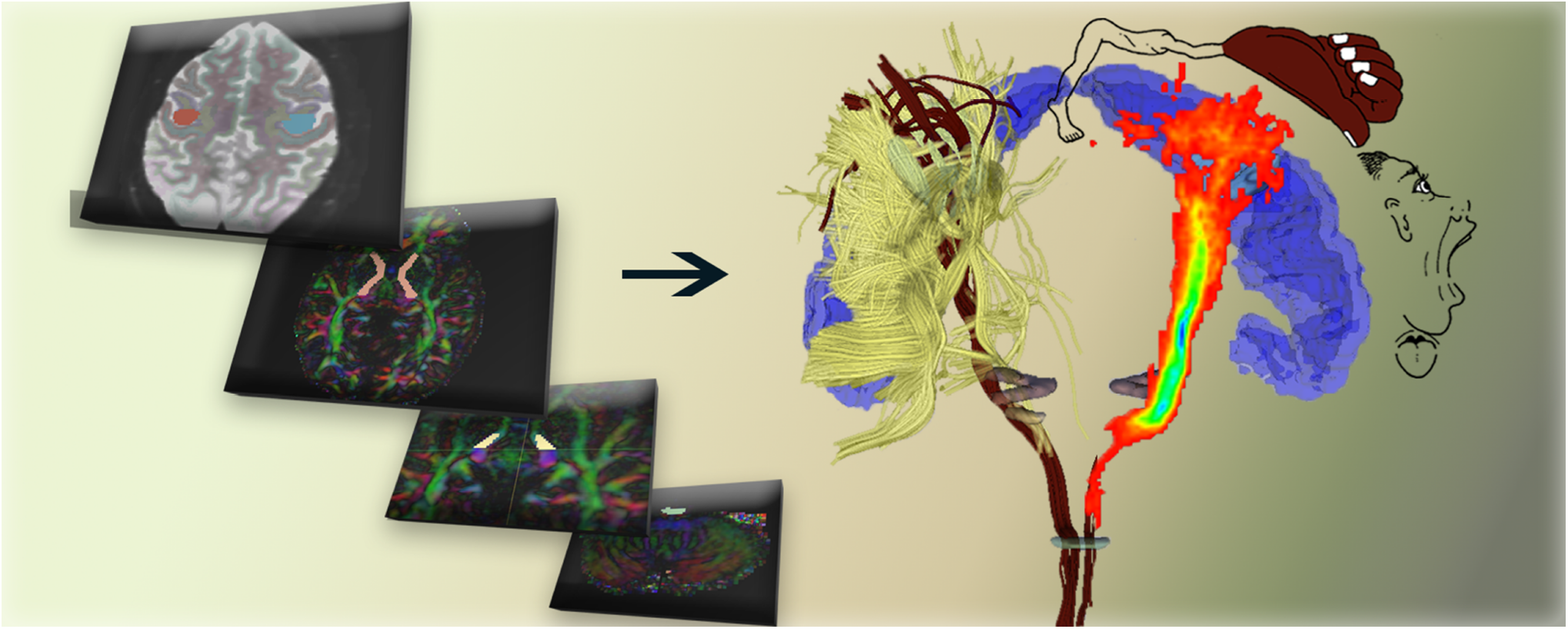

**Highlights:**

- A new method based on three neuroanatomical landmarks using higher-order diffusion MRI tractography for the complete corticospinal tract (CST) delineation is proposed.
- Focus on the clinically significant hand motor area with description of a novel method to isolate and quantify the hand-related motor fiber tracts (HMFTs).
- It is suggested that crossing fibers like the superior longitudinal fascicle are restricting the full representation of the CST in the hand motor area.
- Qualitative and quantitative analysis is performed and variability heatmaps of the human CST and HMFTs are provided based on healthy human subjects from the Human Connectome Project.
- Clinical applications of the CST and the HMFTs tractographic analyses are discussed, along with the historical aspects related to the CST, given its clinical importance for neurology/neurosurgery, neurorehabilitation and regenerative medicine.

## 1 Introduction

Of primary relevance and clinical interest among the fiber pathways in the human brain have been the pathways associated with motor function and, more specifically, the corticospinal tract (CST). Strictly speaking, the CST is the large descending fiber tract that originates in the cerebral cortex and extends longitudinally through the bulbar pyramid. The terms corticospinal tract and pyramidal tract (PT) often are used interchangeably (e.g., (Mai and Paxinos 2011)), and are used synonymously in the present study. The anatomy of the corticospinal tract and its correlation with clinical outcomes has been the focus of medical research since the 1600s. Willis (Willis 1965) described and named the pyramids of the medulla in the macrodissected adult human brain and in other large mammals, stating that they resembled great nerves. Mistichelli (1709, cited in (Finger 1994)) described crossing fibers on the ventral surface of the medulla and published an illustration of the pyramidal decussation. However, he maintained that these fibers originated from the dura mater. One year later, Pourfour du Petit (1710, cited in (Finger 1994), and in (Clarke and O’Malley 1996)) gave a description of the fiber architecture of the pyramidal decussation using postmortem human and experimental animal (dogs) material. He provided his own illustrations of the decussating fibers in the pyramids, and noted that the cerebral cortex has a direct role in movement. It was in the 1800s when Gall and Spurzheim (Gall and Spurzheim 1810) associated the PT with motor function and that Cruveilhier (Cruveilhier 1853) correlated its atrophy with contralateral paralysis. Turck ((Turck 1851); 1852, cited in (Clarke and O’Malley 1996)), using secondary degeneration methods, observed both crossed and uncrossed pyramidal tract fibers. Turck believed the pyramidal tract originated in the basal gray matter of the brain and not in cerebral cortex. Thus, he used "pyramidal" to denote the pyramids of the brainstem rather than Betz cells, which had not yet been described. A few years later, the crossed and uncrossed pyramidal fibers were recognized by Bouchard (Bouchard 1866) as parts of the PT. Finally, the uncrossed lateral PT fibers were identified in the macaque by Schafer (Schafer 1883) as well as in human pathological material by Pitres (Pitres 1884). In 1877, Flechsig (Flechsig 1877), observed in pathological material that lesions of the motor area of the cerebral cortex result in degeneration of the PT. In 1905, Flechsig (Flechsig 1905) noted that the cortical origin of the PT, i.e., the CST, appeared to be the middle and upper thirds of the precentral gyrus. Shortly thereafter, Holmes and May (Holmes and May 1909) provided further insight on the cortical architectonic origins of the CST, identifying the source as the giant pyramidal cells of Betz in Brodmann’s area 4 (BA 4). They performed high cervical hemisections in chimpanzee, monkey, lemur, cat and dog, and observed retrograde changes (reactionary chromatolysis) in Betz cells in Nissl stained serial sections. They conducted postmortem examination of two human patients with high cervical damage and noted similar changes in Betz cells. Since the early 1900s, qualitative studies have been complemented by quantitative investigations (Campbell 1905; Ford and Hackney 1997; Hille 2001; Lassek and Rasmussen 1939; Parent 1996; Pellegrino et al. 1984; Yagishita et al. 1994) that have produced a detailed understanding of the CST in terms of number of cells and fibers composing this fiber tract in humans.

The CST is involved in several complex functions traditionally associated in clinical practice with the mediation of voluntary skilled movements. Experimental studies in the rodent and non-human primate CST have elucidated the anatomy and physiology of the motor system and the CST (Barnard and Woolsey 1956; H.Kuypers 1958; H. G.Kuypers 1964; Levin and Beadford 1938; Nyberg-Hansen and Rinvik 1963; Peele 1942; Russell and DeMyer 1961). In particular, based on some of those studies, the present consensus is that the origin of the CST is more restricted in humans compared to non-human primates. In humans the CST is currently thought to originate from BA 4 (primary motor cortex, giving rise to about 60% of the CST fibers), area 6 (supplementary motor areas) and part of the parietal lobes (Davidoff 1990). These studies are highly relevant, given their translational potential to humans, especially in the clinical domain of neuroregeneration and repair. Importantly, in order to be able to screen for pathological findings, track clinical progress and detect the regenerative potential of tested therapies, scientists need to know the anatomy of the individual subject.

The advent of diffusion magnetic resonance imaging (dMRI) (Basser et al. 1994; Le Bihan et al. 1986) has greatly facilitated the study of neural connections in humans, both ex-vivo and, most importantly, in-vivo in a non-invasive fashion. Although dMRI is a powerful approach for studying brain connections there are limitations that need to be overcome. Both the diffusion tensor imaging (DTI) and high angular resolution diffusion imaging (HARDI) techniques (Basser et al. 1994; Tuch et al. 2002) have permitted a more valid examination of the most sizeable and best-defined fiber tracts in humans (Nikos Makris et al. 2005a; Schmahmann and Pandya 2006). Thus, pioneering studies using diffusion imaging-based segmentation (N. Makris et al. 1997; Nikos Makris et al. 2005a) and tractographic approaches (Lori et al. 2002; Mori et al. 1999) aimed at delineating the corpus callosum (Huang et al. 2005), CST (Stieltjes et al. 2001), optic radiations (Sherbondy et al. 2008) and the major corticocortical association fiber tracts (N.Makris et al. 1997, p. 199;Mori et al. 1999) in humans. Given the anatomical location of the hand representation in the posterior limb of the internal capsule (IC) (Bertrand et al. 1965; Chronister and Hardy 1997) and its considerable size and well known anatomy in humans, the CST may be one of the most highly studied fiber pathways in the human brain (Betz 1874; Lassek and Rasmussen 1939; Parent 1996). Several investigations of its physiology (Schäfer 1910), function (Bertrand et al. 1965), clinical significance (Dejerine and Dejerine-Klumpke 1895; Hirayama et al. 1962), and MRI-based characterization (Ellis et al. 2001; Holodny et al. 2005; Pierpaoli et al. 2001; Yagishita et al. 1994) have yielded information indicating the importance of accurately and precisely mapping its different components, and the hand area in particular, given its key clinical significance for motor and sensory functions in humans (Brothwell 1960; Skirven et al. 2011). Nevertheless, despite the great clinical significance of the hand motor area for a patient’s independence in terms of the activities of daily living, the hand-related motor fiber tracts (HMFTs) have not been explored as part of the CST using solely dMRI tractography. Therefore, an assessment of the anatomical variability of the CST and HMFTs is critical. It is essential that such studies become a bridge between human brain anatomy and function, to allow for a more reliable prediction of the neuroregenerative impact of novel therapies on human subjects, utilizing non-invasive imaging tools with objective scales. Traumas or diseases of the CST lead to significant restriction of human locomotion. The CST is affected in several neurological entities, such as stroke and spinal cord injury (SCI) and several neurodegenerative diseases (Ropper et al. 2014). Stroke and SCI alone have a tremendous impact on the general population with huge personal and financial costs to society. Thus clinical trials involving molecules, cells or even biomaterials, aimed at minimizing the impact of CST trauma or even reversing it, creating an appropriate environment for regeneration and repair are currently underway involving interdisciplinary research. MRI diffusion tensor imaging, and especially tractography, is an excellent tool for the establishment of the grounds for a common scale “language” among interdisciplinary teams. dMRI tractography could pave the way for development of an objective scale and could catalyze the discovery of new interventions that could benefit a vast population of CST patients.

Neurosurgical candidates with tumors in proximity to the CST are highly prone to the collateral effects of tumor excision and the possibility of being left paralyzed in an attempt by the surgeon to save their lives. Currently, there is no single method for tracking the CST for surgical planning with accuracy and reliability. Moreover, beyond treatment purposes, understanding the causal role of the CST for specific neurological conditions is also of great importance. Several genetic studies are trying to correlate the genotype with the phenotype of several neurologic diseases, such as ALS, in order to elucidate causes of the disease (Ravits et al. 2013). Unfortunately, only pathological data from postmortem studies are available for exploration in the human population. Thus, pursuing non-invasive in vivo imaging of the the CST using dMRI is an attractive prospect from both clinical and basic research perspectives. Diffusion MRI tractography has been widely used in pre-surgical planning, with the CST being one of the tracts that has been widely studied due to its importance for voluntary movement (Berman et al. 2004; Bucci et al. 2013; Chen et al. 2016a; Kinoshita et al. 2005; Snow et al. 2016).

Given the great clinical importance of the HMFTs within the CST, we designed experiments to test our hypothesis that dMRI tractography can be used to adequately delineate the HMFTs as part of the CST. This delineation allows for differentiation from the dense surrounding tracts in that region, namely the arcuate fascicle (AF), superior longitudinal fascicles (SLFI, SLFII, SLFIII) and subdivisions of the corpus callosum (CC). Furthermore, we assessed the anatomical variability of the CST in its entirety and the HMFTs in particular. We performed 2-tensor (2T) tractography in 37 subjects from the Human Connectome Project (HCP). Using MRI identifiable, anatomically robust ROIs, namely the pyramids, cerebral peduncles and internal capsule (Nikos Makris et al. 1999; Parent 1996) we were able to delineate the CST in its entirety as well as to isolate its hand motor representation. Furthermore, we estimated their variability and generated a database of their volume, length and biophysical parameters. We have used unscented Kalman filter based 2T tractography (Malcolm et al. 2010), which was one of the winners of the fiber cup competition (Fillard et al. 2011). This algorithm can robustly trace fibers through crossing fiber regions. To the best of our knowledge, this is the first attempt to fully delineate the HMFTs as part of the CST using dMRI tractography, including determining the variation of the tracts within the healthy population. To the best of our knowledge, this is the first attempt to isolate and quantify the hand motor area using purely dMRI tractography.

## 2 Methods

### 2.1 Aims

We used dMRI two-tensor unscented Kalman filter deterministic tractography (UKF deterministic tractography) in 37 healthy human subjects with state-of-the-art data to accomplish four main goals: a) to anatomically delineate the complete corticospinal tract, b) to isolate the hand-related motor area as a subset of the CST, c) to generate a database of volume, length and biophysical parameters such as fractional anisotropy (FA) (Basser 2004), axial diffusivity (AD) (Song et al. 2003), and radial diffusivity (RD) (Song et al. 2002, 2003) for the CST and for the HMFTs, and d) to quantify the variability of the CST and of the HMFTs within the healthy population using probabilistic variability maps (heat map).

### 2.2 Subjects

Thirty-seven healthy adult subjects (mean age=30.88, SD=3.14), 19 male subjects (mean age=30.42, SD=3.37) and 18 female subjects (mean age=31.36, SD=2.89) were used for this analysis from the state-of-the-art WashU Connectome Project led by Washington University (http://www.humanconnectome.org/data/), University of Minnesota and Oxford University (the WU-Minn HCP consortium). Almost 65% of the tested subjects were 31-35 years old (45.8% male subjects and 54.2% female subjects, within that age group). Of the total subjects, 27% were 26-30 years old (60% male subjects and 40% female subjects, within that age group) and only 8% were 22-25 years old (66.7% male subjects and 33.3% female subjects, within that age group).

### 2.3 MRI procedures

The data were acquired with the specially configured WU-Minn Skyra Connectome scanner from Siemens using a 32 channel standard Skyra coil. Customized hardware with gradient coil and gradient power amplifiers for research use increased the maximum gradient strength from standard 40 mT/m to 100 mT/m on the WU-Minn 3T scanner. Diffusion data sets were acquired with 1.25 × 1.25 × 1.25 mm^3^ voxels, FOV = 210mm, 111 slices, TE = 89.50ms, TR = 5520ms, b-value= 1000, 2000, 3000s/mm2, 90 directions per b-value and 18 non-diffusion-weighted volumes (Uğurbil et al. 2013). For our analysis we used only the b-value 3000s/mm^2^ data.

### 2.4 Tractographic Delineation of the Corticospinal Tract

This dataset is a subset of the publicly available Human Connectome Project (http://www.humanconnectome.org/data/). Therefore, it was already processed through the Human Connectome Project preprocessing pipeline described in the original paper (Glasser et al. 2013). UKF tractography was performed using a multi-tensor tractography algorithm (Malcolm et al. 2010). Based on prior results, two-tensor UKF tractography (Malcolm et al. 2010) was chosen as a more sensitive method for the delineation of the CST, allowing fiber tracing in areas that are known to be heavily affected by crossing fibers and branching (Baumgartner et al. 2012; Chen et al. 2016a). In an attempt to increase accuracy, aiming at future clinical applicability of the method, we chose to manually draw certain regions of interest (ROIs) using anatomical landmarks for the CST and hand motor area rather than the automatic methods. We then used the White Matter Query Language (WMQL) to delineate the fiber tracts passing through all the drawn ROIs and then proceeded with the quantification methodology.

#### 2.4.1 Two-tensor Deterministic Tractography

After completing the quality control for the UKF tractography, the 3D Slicer software package (version 4.4.0, www.slicer.org) via the SlicerDMRI (Fedorov et al. 2012; Norton et al. 2017) project (dmri.slicer.org) was used to create the tensors mask and then sample the ROIs. The CST was delineated using three ROIs, which were manually-drawn in 3D Slicer in axial color-coded dMRI sections as shown in detail in Fig. 1. Specifically, the first ROI was in the upper medulla rostral to the inferior olivary nucleus (hereinafter referred to as medulla-ROI), the second ROI was at a mesencephalic level rostral to the substantia nigra and red nucleus (hereinafter referred to as brainstem-ROI) and the third ROI was in the internal capsule (IC) at the anterior commissure (AC) level (hereinafter referred to as capsular-ROI) (Fig. 1). In the medulla- and brainstem-ROIs we selected only voxels with fibers in the vertical dimension (Z-axis), which are color-coded blue. In the capsular-ROI we included all voxels of the IC.

**Fig. 1.**
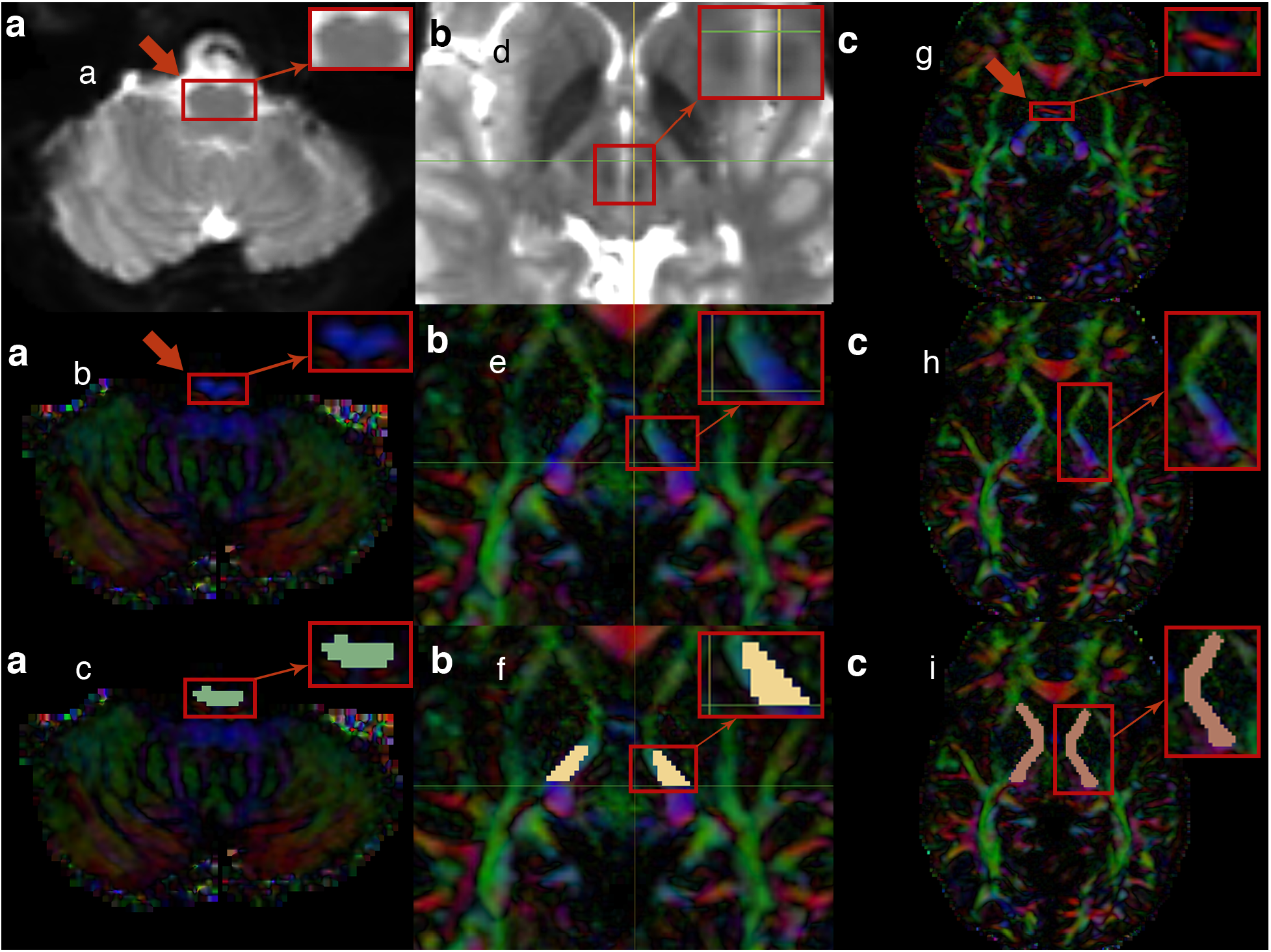
***a (left portion)*** The upper medulla oblongata, rostral to the inferior olivary nucleus, was located in the baseline (b0) MRI brain images (axial plane) as indicated by the arrow (**a**). The same area is illustrated in the diffusion tensor imaging (DTI) (axial plane) as indicated by the arrow (**b**). Finally, the medulla-region-of-interest (medulla-ROI) was drawn to the targeted region (axial plane) (**c**). ***b (middle portion)*** The region included in the square was located in the baseline (b0) MRI brain images in order to detect the red nucleus (axial plane) (**d**). The region of the cerebral peduncle of the brainstem rostral to the substantia nigra and red nucleus (thereby, rostral to the horizontal level marked by the green line in the image) was targeted (axial plane) (**e**). Finally, the brainstem-ROI was drawn to that area (axial plane) (**f**). ***c (right portion)*** The anterior commissure (AC) level was located as indicated by the red color-coded fibers that disappear in the next image (axial plane) (see the red arrow) (**g**). Just after the AC was no longer visualized in the DTI and moving superiorly, the internal capsule (IC) was targeted as illustrated (axial plane) (**h**). Therefore, the capsular-ROI was drawn to the IC at the level of the AC (axial plane) (**i**).

WMQL software (Wassermann et al. 2016) was used for the extraction of fiber bundles based on the manually drawn ROIs so that the fibers passed through all three ROIs. Subsequently, the results were visualized in 3D Slicer for quality control and refined by removing any commissural fibers that were not part of the CST.

### 2.5 Quantitative Analyses

The tract biophysical characteristics (i.e., tract volume, tract length, FA, RD, AD) were obtained in all 37 healthy human subjects. The diffusion imaging biophysical parameters of FA, AD and RD may relate to fiber tract coherence and integrity (Basser 2004; Song et al. 2002, 2003). We normalized the tract volume results by the whole brain volume for each individual to avoid any inconsistencies due to personal characteristics.

Symmetry index (SI) 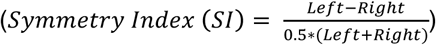 for left and right tracts (Galaburda et al. 1987) for measures of volume, mean FA, mean AD and mean RD were calculated for each individual, and across subjects.

Also, the presence of outliers in the above-mentioned measures such as FA, AD, RD and volume was tested using Tukey’s range test in the initially delineated CSTs and then in the delineated HMFTs (Tukey 1977).

### 2.6 Intra-rater and inter-rater reliability

The two raters (K.D. and M.T.) received independent training and worked independently during the rating stage to avoid any bias.

For the inter-rater reliability process, K.D. processed all 37 WashU subjects and M.T. processed 10 randomly selected WashU subjects independently. The results for the 10 common cases were compared for the two independent raters using the Cronbach’s Alpha reliability analysis in SPSS v23 statistical software package (Cronbach 1951).

To assess intra-rater reliability, M.T. processed the previously randomly selected 10 WashU cases for a second time several days after the initial processing, without referring to the previous results. For this process the results for those 10 cases were compared using the SPSS v23 statistical software package. As for the inter-rater reliability, the intra-rater reliability assessment was done by performing the Cronbach’s Alpha reliability analysis.

### 2.7 Isolation of the hand-related motor fiber tracts

Based on the previous work of Yousry et al. (Yousry et al. 1997) who used fMRI to accurately define the hand motor area in structural MRI scans, the hand motor area was defined in the axial plane as a knob-like, broad based, posterolaterally directed region of the precentral gyrus. This region usually has an inverted omega shape and sometimes a horizontal epsilon shape, with a mean diameter of 1.4 cm. On average it is located about 23 mm from the midline, just posterior to the junction of the superior frontal sulcus with the precentral sulcus and 19 mm from the lateral surface. In the sagittal plane, this knob takes the form of a posteriorly directed hook with a mean depth and height of 17 and 19 mm, respectively. It is located in the sagittal plane of the same section in which the insula can be identified, perpendicular to its posterior end. In neurologically unaffected hemispheres like the ones we used in the present study, the sensitivity of detecting the hand motor area in a structural MRI scan with this method has been shown to be 97–100%, with an accuracy of 97–100% (Yousry et al. 1997).

To isolate the hand motor area, we used 3D Slicer to draw a fourth ROI in the area of the “omega sign” in the axial plane, and the drawing process was completed in the coronal plane (hereinafter referred to as cortical-ROI). In order to ensure the successful location of the “omega sign” in the precentral gyrus we used in parallel the FreeSurfer labelmaps created by the process of automatic subcortical segmentation for each subject. By using the slice intersection feature of 3D Slicer the same area was located in the coronal plane. The remaining non-colored area of gyri, defined by the previously drawn colored marks of the axial plane, was filled in with the same color, as illustrated in Fig. 2a. All four ROIs (the previously drawn three ROIs and the new one for the hand area) were merged into one labelmap, which was then used with the WMQL software to extract the fiber bundles emanating from all four ROIs, thereby isolating the HMFTs (Fig. 2b). The isolated HMFTs represented a subset of the initial CST and required no further refinement. We note that using this manual procedure to select the ROIs makes the delineation of the fiber tracts robust to errors in EPI distortions in the brain stem region as reported in Irfanoglu et al. (Irfanoglu et al. 2015).

**Fig. 2.**
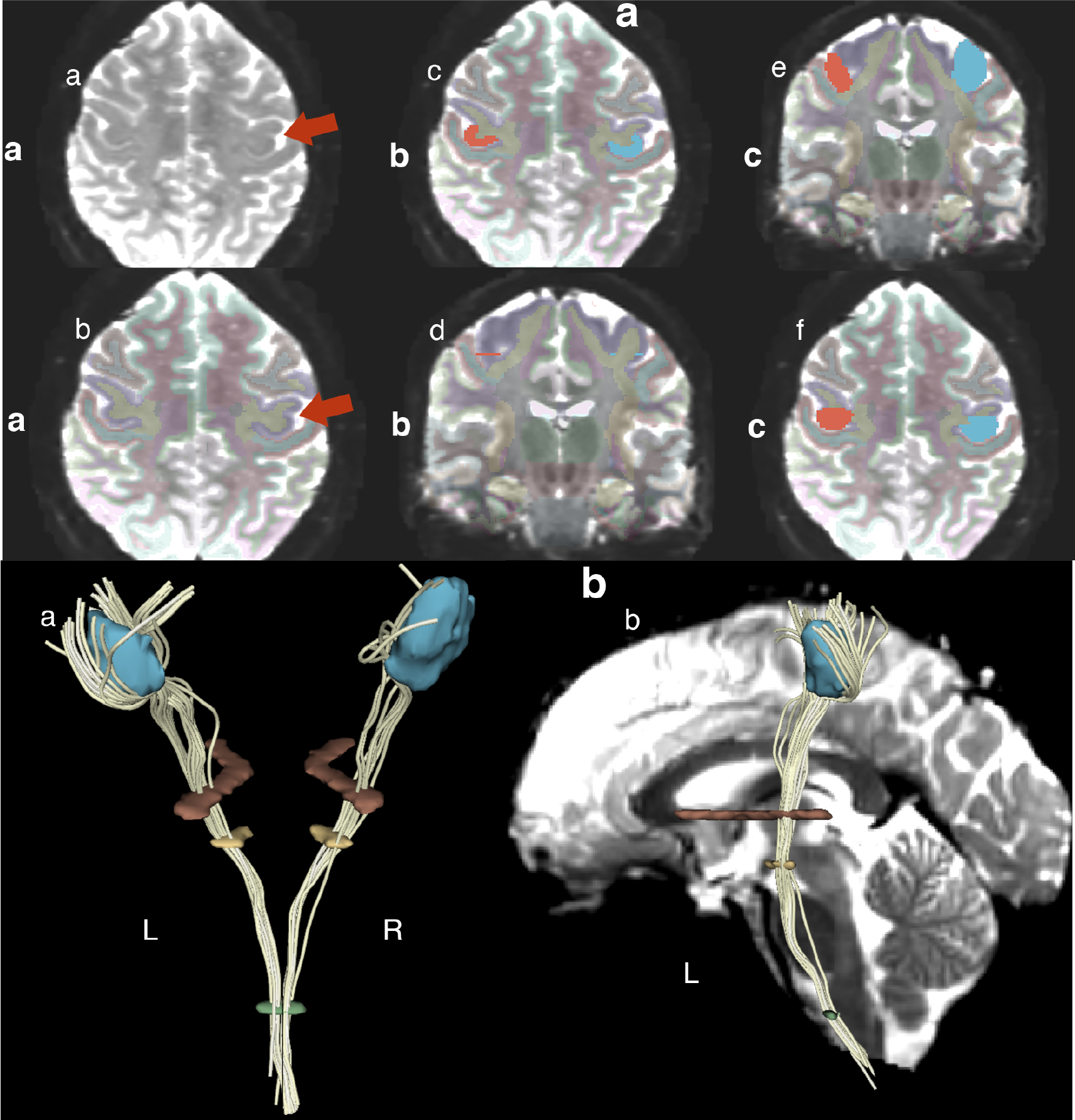
***a (upper portion)*** Illustrative demonstration of the hand motor area isolation methodology using a fourth region of interest (ROI) in the precentral gyrus (cortical-ROI). ***a (sub-section on the left)*** Detection of the “omega sign” in the precentral gyrus as indicated by the arrow (axial plane) (**a**). Confirmation of the correct localization via overlaying the FreeSurfer labelmap for each case that was a result of the automatic FreeSurfer subcortical segmentation. The purple color indicates the precentral gyrus (axial plane) (see the red arrow) (**b**). ***b (sub-section in the middle)*** Drawing of the cortical-ROI in the cortical region of the “omega sign” (axial plane) (**c**). Detection of the same area in the coronal plane utilizing the “slice intersections” tool of 3D Slicer (coronal plane). Note that either a straight line covering the entire extent of the gyrus of interest or two bullets defining the gyrus of interest can be observed as illustrated in the image (**d**). ***c (sub-section on the right)*** The remaining non-colored area of the gyrus of interest was filled in with the same color to complete the cortical-ROI (coronal plane) (**e**). In the axial plane, the correct area of interest, namely the hand motor area, was verified (**f**). ***b (lower portion)*** Spatial representation of all four ROIs after they were added to one labelmap in order to isolate the hand-related motor fiber tracts (HMFTs). HMFTs were defined in White Matter Query Language (WMQL) as the fiber tracts passing through all four ROIs, as demonstrated in the case of that HMFT. Posterior view of a 3D model of the four ROIs and the HMFT passing through all four in 3D Slicer software. Left and right sides are designated L and R, respectively (**a**). Lateral view (left side) of the same subject’s HMFT with an added sagittal MRI section of the subject’s b0 MRI image to illustrate spatial localization of the ROIs (**b**)

### 2.8 Assessment of inter-subject variability of CST and HMFTs

We generated probabilistic maps for the CST and the HMFTs in Montreal Neurological Institute (MNI) 152 standard space, by first registering each subject to an MNI152 template with a resolution of 1mm^3^. Affine transformation was used for the co-registration of our fiber bundles and the baseline b0 brain MRI images. The CSTs and the HMFTs were processed with the pipeline described above.

We calculated a fiber tract mask on the MNI152 template for each subject using 3D Slicer software. In this mask a voxel had a value only if a tract traversed it. Finally, we averaged the masks for all our subjects in order to produce the probabilistic heat map for the CST and repeated the same process for the HMFTs.

### 2.9 Crossing fibers interference analysis on HMFTs

We ran two sets of experiments to determine the effect that crossing fibers at the level of the CST have on our hand motor area findings.

The first set of experiments entailed the delineation of the AF and the superior longitudinal fascicles, i.e., SLF I, SLF II and SLF III using the WMQL software’s automatic fiber extraction method (http://tract-querier.readthedocs.io/en/latest/#) based on previously described definitions of the tracts (Wassermann et al. 2013). The second set of experiments involved the delineation of the CC fiber tracts based on a variant of the WMQL-based published methodology (Wassermann et al. 2016) to isolate the 7 sub-sections of the CC. The quantification and the MNI space co-registration were performed as described in sections 2.5 and 2.8, respectively.

## 3 Results

### 3.1 Delineation of the CST and HMFTs

This study demonstrated clear delineation of the CST using state-of-the-art data, with the combination of both the tractographic algorithm and 3 manually drawn ROIs. Fig. 3 illustrates a representative example of a CST delineation using our methodology. Our approach allowed us to delineate the CST without fiber truncations and with a limited effect of crossing fibers interference. Nevertheless, as indicated by Fig. 4, the hand motor area, which is the focus of our study due to its clinical significance, still seems to be affected by technique-related limitations. Fig. 4 is a representative example, demonstrating a significant paucity of fibers to the hand motor area. It should be noted that in our sample of 37 subjects, there was only one case showing extremely poor CST representation and, therefore, was considered an outlier and was not included in the HMFT analysis.

**Fig. 3.**
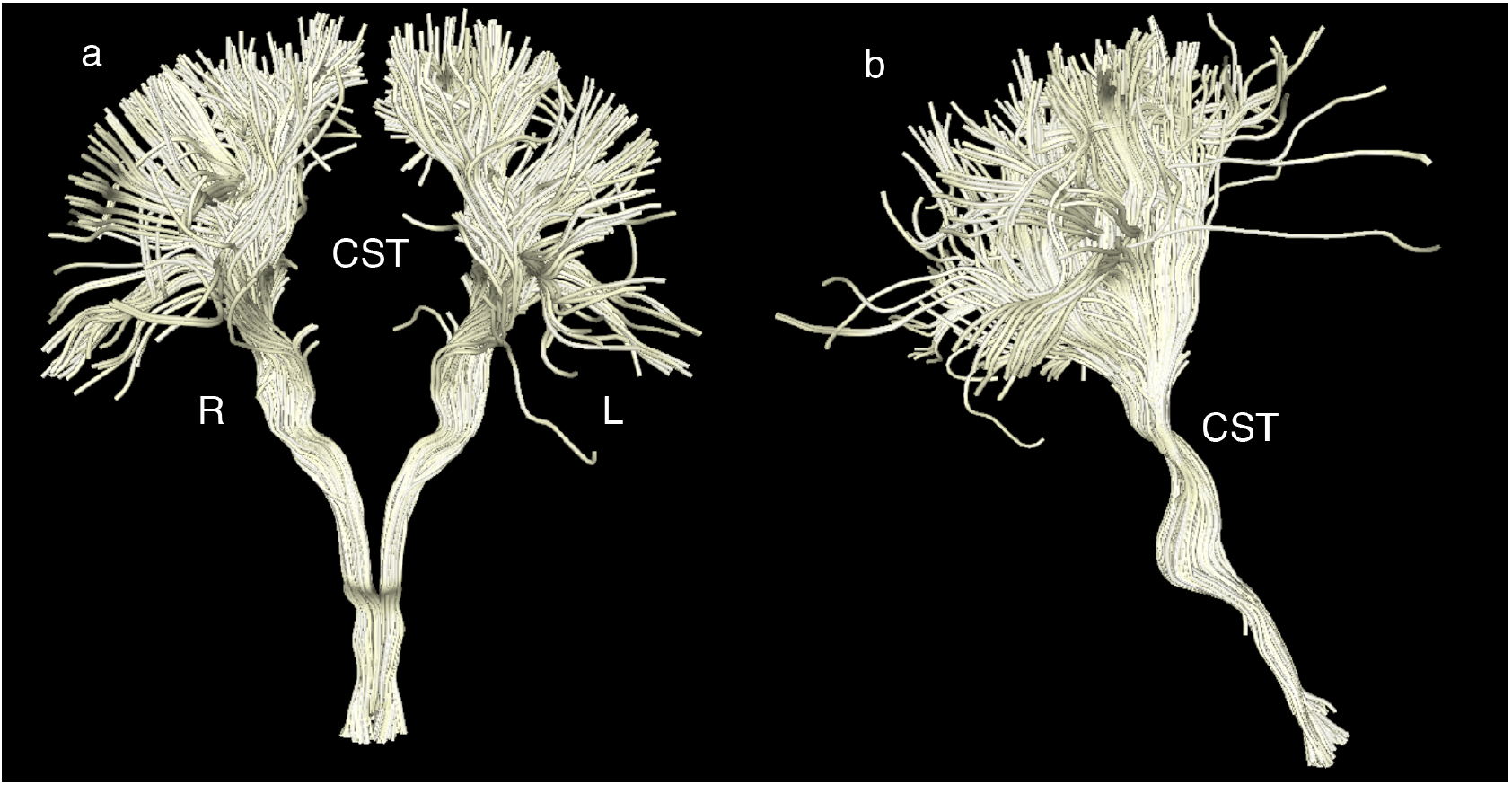
Representative corticospinal tract (CST) delineation using the three manually drawn regions of interest (ROIs) (in the medulla, brainstem and internal capsule as described in section 2.4.1) for maximum precision. **a** Front view of the CST in the coronal plane. The left side is labeled L and the right side R. **b** Side view (right side) of the CST (coronal plane)

**Fig. 4.**
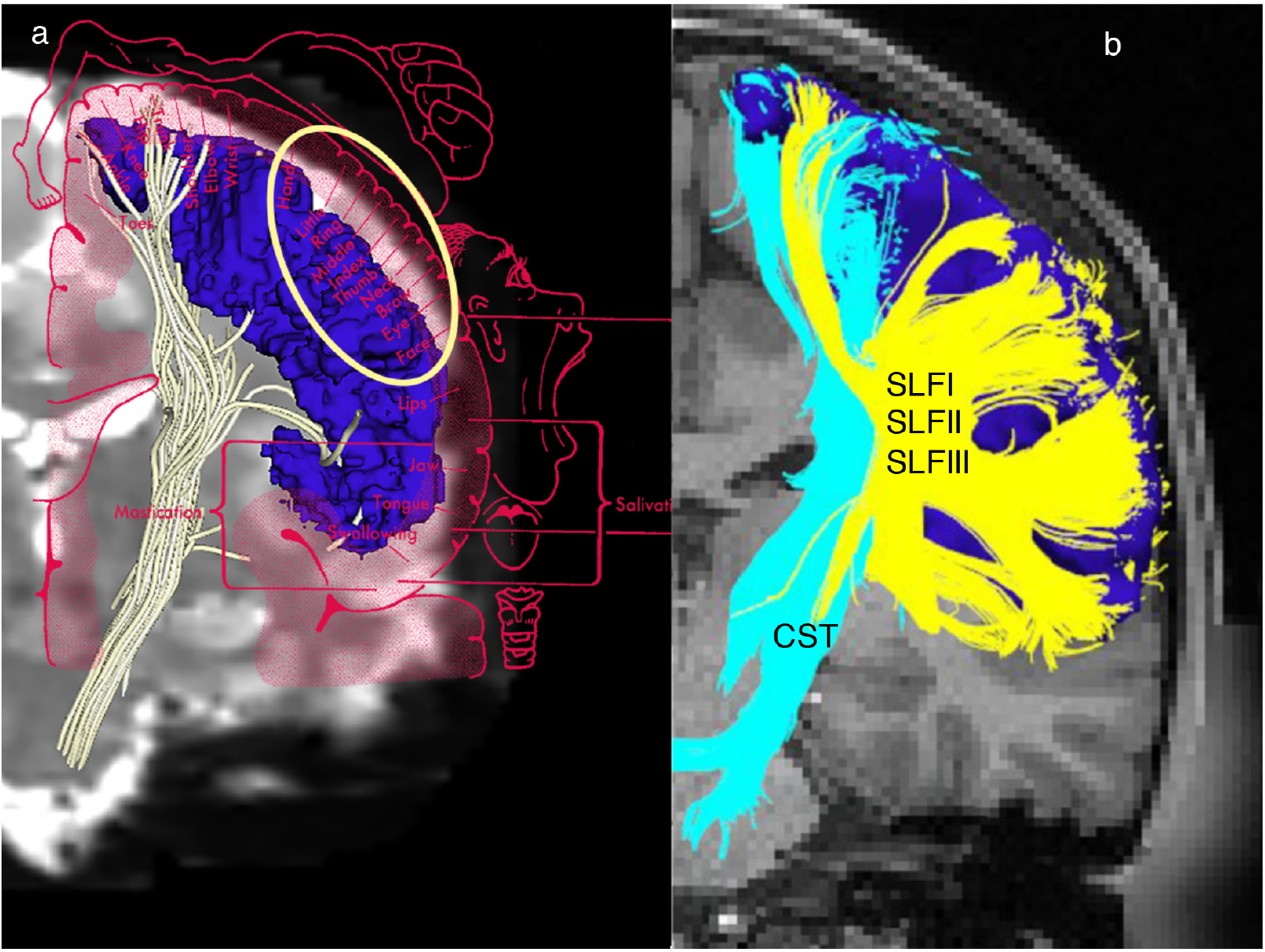
**a** A representative corticospinal tract (CST) of a healthy human subject delineated by diffusion magnetic resonance imaging (dMRI) tractography using high-resolution data of the HCP (Washington University; abbreviated as WashU) sample. Please note the paucity of fibers in the lateral aspect of the hemispheres. This is possibly due to limitations of dMRI tractography. When overlaid with the motor homunculus (coronal plane), the lateral part of the motor homunculus indicates that the hand motor area, which should have been a part of the CST, is significantly limited, something that can possibly be explained by the presence of massive superior longitudinal fascicles I, II and III (SLF I, II and III) crossing fibers at the centrum ovale. **b** In this representative case, the entire precentral cortex (dark blue) has been used as the seed to grow the CST (light blue). Crossing fibers at the centrum ovale relating principally to the superior longitudinal fascicles I, II and III (SLF I, SLF II and SLF III, in yellow) (Nikos Makris et al. 2005b) do not allow effective sampling of CST from the lateral aspect of the precentral gyrus (motor cortex)

Upon successful delineation of the CST, the hand motor area was isolated as described in section 2.7, adding the cortical-ROI in our method. Despite existing limitations, the HMFTs were successfully isolated in 36 of 37 cases. The only case with an absent HMFT was that with the initially poor CST; therefore, this result was not associated with the HMFTs-related methodology. As expected from prior experience and our CST results, the hand motor area isolation was much more challenging and the paucity of fibers in that region was clear in some subjects. Moreover, a greater inter-subject variability was noted on the initial visual inspection of the HMFTs. A representative example of our isolated HMFTs is shown in Fig. 5.

**Fig. 5.**
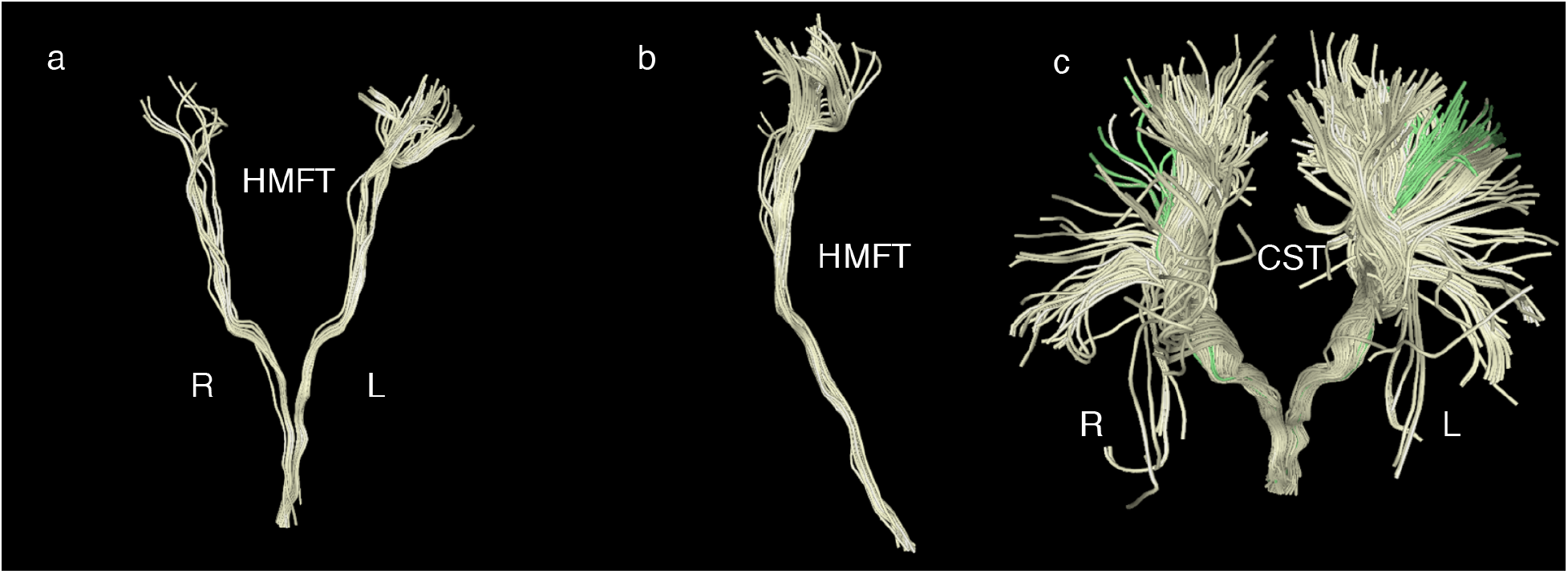
Representative hand-related motor fiber tracts (HMFTs) delineation using the four manually drawn regions of interest (ROIs) (in the medulla, brainstem, internal capsule and the “omega sign” in the precentral gyrus, as described in section 2.7) for maximum precision. It should be noted that, likely due to the current limitations of diffusion resonance imaging (dMRI) tractography, we could not obtain an optimal representation of the hand motor area in all subjects and there was significant variability among the subjects, contrary to our corticospinal tract (CST) results. **a** Front view of the HMFTs in the coronal plane. The left side is labeled as L and the right as R. **b** HMFTs viewed from the right side (coronal plane). **c** Front view of the CST (light yellow). The HMFT (green) is depicted as a subset of the CST for direct comparison

### 3.2 Intra-rater and inter-rater reliability

Cronbach’s Alpha values were used to determine inter- and intra-rater reliability for the present technique, which was found to be highly replicable. The most reliable values in our study are for the diffusion parameters (i.e., FA, RD, AD values), with Cronbach’s alpha approaching the maximum of 1.00 (0.95 for FA inter- and FA intra-rater reliability; 0.99 for RD inter- and intra-rater reliability; 0.98 for AD inter- and intra-rater reliability). Tract volume was found to be moderately reliable (0.80 for inter-rater and 0.90 for intra-rater reliability) and the number of fibers least reliable (0.70 for inter-rater and 0.68 for intra-rater reliability), in agreement with previous studies (Carlson et al. 2014; Dini et al. 2013).

### 3.3 Quantitative Analyses

We generated a database of biophysical parameters (i.e., FA, AD, RD) and volume of the CST and the HMFTs (Tables 1 and 2).

**Table 1.**
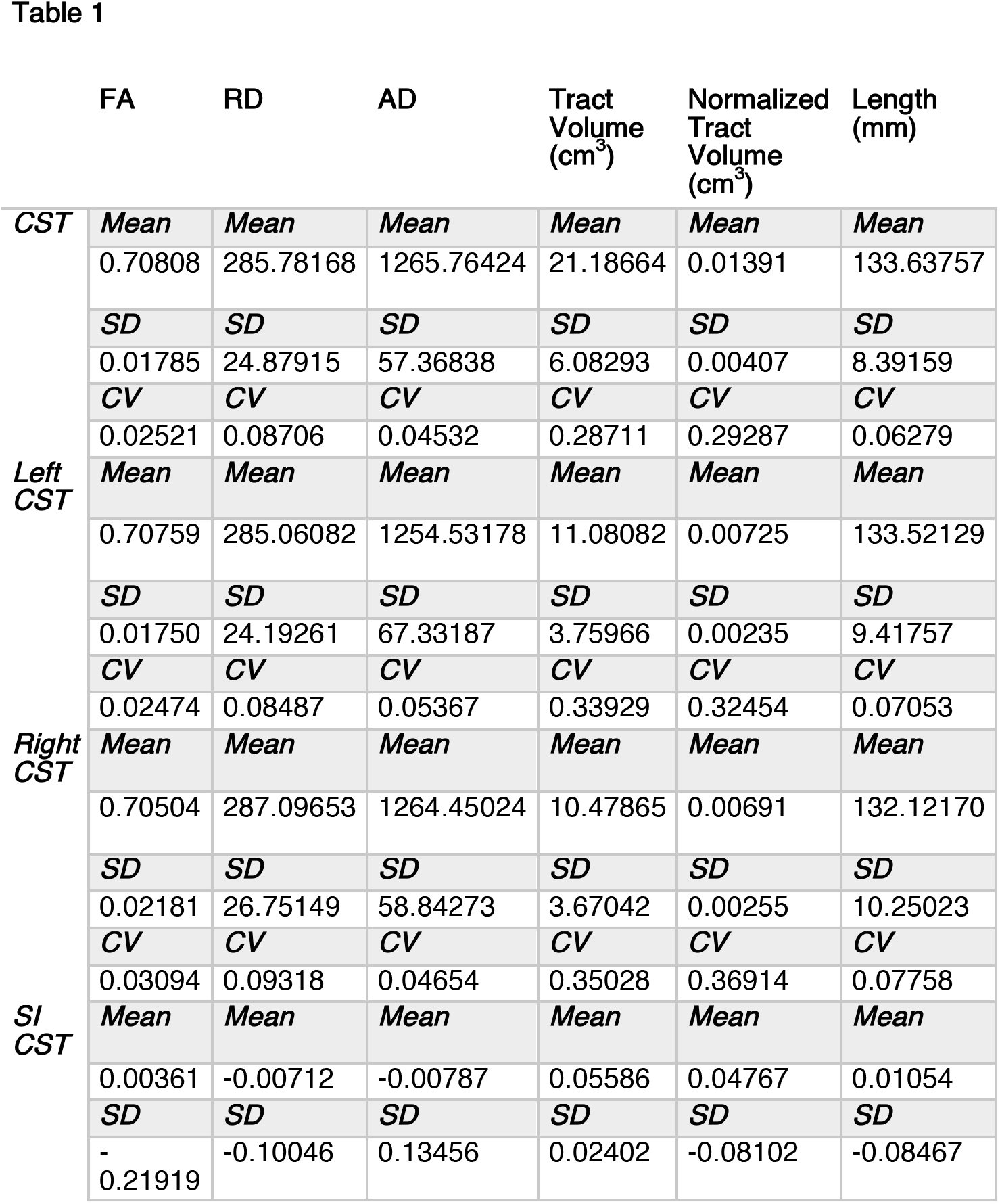
Summary of the quantitative analysis of the biophysical characteristics and tract volumes of the delineated CSTs. Note that the coefficient of variation is very small (<10%), indicating minor variability of the results among our healthy subjects. Tract volume, by contrast, was the only parameter that was highly variable among the subjects.

**Table 2.**
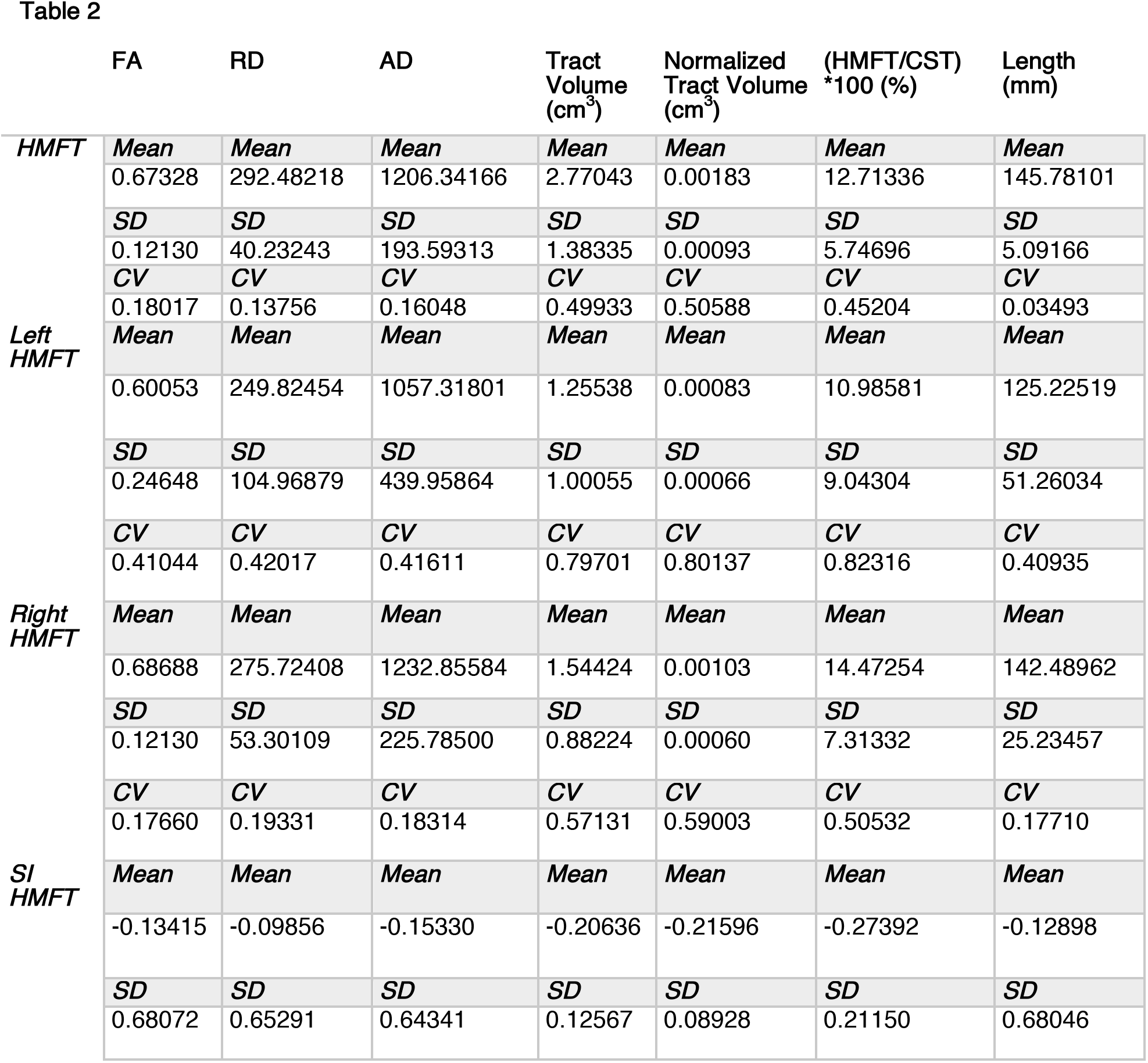
Summary of the quantitative analysis of the biophysical characteristics and tract volumes of the delineated HMFTs. Note the increase in the coefficient of variation compared to the CST measurements. This indicates a much higher variability of the results among our healthy subjects (>10%), possibly due to the limitations of the currently available imaging tools, leading to outliers. In agreement with the CST quantitative measurements database, the tract volume was found once again to be much more variable among the subjects (coefficient of variation approximately 53%). Note that 12.7%, SD=5.7, of the CSTs analyzed refers to the hand motor area. This measurement was based on the normalized volumes of the respective tracts.

The CST had a mean volume of 21.187 cm^3^, SD=6.083 (left CST: mean volume=11.081 cm^3^, SD=3.760; right CST: mean volume=10.479 cm^3^, SD=3.670). After normalizing the tract volumes by dividing by the total brain volume, to give a fraction of the whole brain volume, the CST mean volume fraction was 0.014, SD= 0.004 (left CST: mean volume fraction= 0.007, SD= 0.002; right CST: mean volume fraction= 0.007, SD= 0.003). The mean FA value of the CST was 0.708, SD=0.018 (left CST: mean FA= 0.708, SD=0.018; right CST: mean FA= 0.705, SD=0.022). The coefficient of variation was <10% for all parameters tested, except for the tract volume which was highly variable between the subjects (coefficient of variation approximately 30%). Overall, FA was the most reliable parameter for our study with a coefficient of variation of only about 2.5%. This agrees with other reports noting the same variability trend (Carlson et al. 2014).

Regarding the HMFTs, the mean volume was 2.770 cm^3^, SD=1.383 (left HMFTs: mean volume=1.255 cm^3^, SD=1.001; right HMFTs: mean volume=1.544 cm^3^, SD=0.882). After normalizing the tract volumes by dividing by the total brain volume, to give a fraction of the whole brain volume, the HMFTs mean volume fraction was 0.002, SD= 0.001 (left HMFTs: mean volume fraction= 0.001, SD= 0.001; right HMFTs: mean volume fraction= 0.001, SD= 0.001). The mean FA values of the HMFTs was 0.673, SD= 0.121 (left HMFTs: mean FA= 0.601, SD= 0.246; right HMFTs: mean FA= 0.687, SD= 0.121). The coefficient of variation was much higher for the HMFTs for all the parameters tested and especially for the tract volume, which was the most highly variable parameter among the subjects (coefficient of variation >50%).

A factor that could contribute to the difference in the coefficient of variation is the presence of outliers. Based on Tukey’s range test, only one outlier was found in the quantitative analysis of the CSTs, which was also in agreement with visual inspection. Thus, we conducted our HMFTs-related quantitative analysis with all 37 subjects and then re-ran it after excluding only that one subject. The quantitative values for the HMFTs mentioned in section 3.3., including the values depicted in Table 2, refer to the 36 subjects’ measurements, given the problematic CST of the excluded subject. Contrary to our CST results that included only 1 outlier, 8 additional outliers were detected within the new total of 36 subjects in our HMFTs analysis. All the outliers were based exclusively on the basis of significant differences in the FA values. Therefore, according on our results, the FA seems to be the most sensitive parameter tested. The presence of outliers leads to the higher variation noted in the HMFTs measurements.

Another finding that could provide an indication of the current capability of isolating and quantifying the hand motor area for clinical purposes regards the percentage of CST fibers belonging to the hand motor area. Based on our analysis, 12.7%, SD=5.7 of the CST fibers were HMFTs (see “(HMFT/CST) * 100 (%)” results reported in Table 2) with the same percentage found to be 11.0%, SD=9.0 for the left side and 14.5%, SD=7.3 for the right side. Once again, consistent with the high variation in determining the hand motor area using dMRI tractography, these percentages demonstrate high variance.

#### 3.3.1 Analysis of symmetry

Tables 1 and 2 list the SI for the left and right measurements of tract volume, length, mean FA, mean AD, mean RD in the CST and HMFTs, respectively.

For HMFTs analysis, given that FA values and left HMFTs volume values did not satisfy the normality assumption, a non-parametric related-samples Wilcoxon Signed Rank Test was run in SPSS, which indicated a significant difference between the FA values of the left and right hand motor region (p<0.05). In particular, the median FA values for the left HMFTs, median=0.69693, were significantly lower than the median FA values for the right HMFTs, median=0.71146, Z=0.486, p=0.016 < 0.05. Six of the outliers detected in our HMFTs isolation were related to asymmetrical values of FA. In particular, an absence of fibers was noted on the left side of five of the aforementioned outliers. Only one of those outliers was lacking fibers to the right side. Otherwise, no statistically significant asymmetry was found in our results.

In terms of the CST analysis, the normality assumption was satisfied, therefore we conducted a parametric paired t-test in SPSS. The test indicated that there was no statistically significant asymmetry in the delineated CST (p>0.05 for all pairs tested).

### 3.4 Assessment of inter-subject variability of CST and HMFTs

We generated probability maps for the CSTs and HMFTs as shown in Fig. 6, following co-registration to the standard MNI152 space. The CST inter-subject variability was significantly lower in the areas adjacent to the drawn ROIs, with higher variability for the cortical areas. The inter-subject variability of the HMFTs, by contrast, seemed to be higher overall. In agreement with the CST results, cortical areas of the HMFTs were much more variable among healthy individuals.

**Fig. 6.**
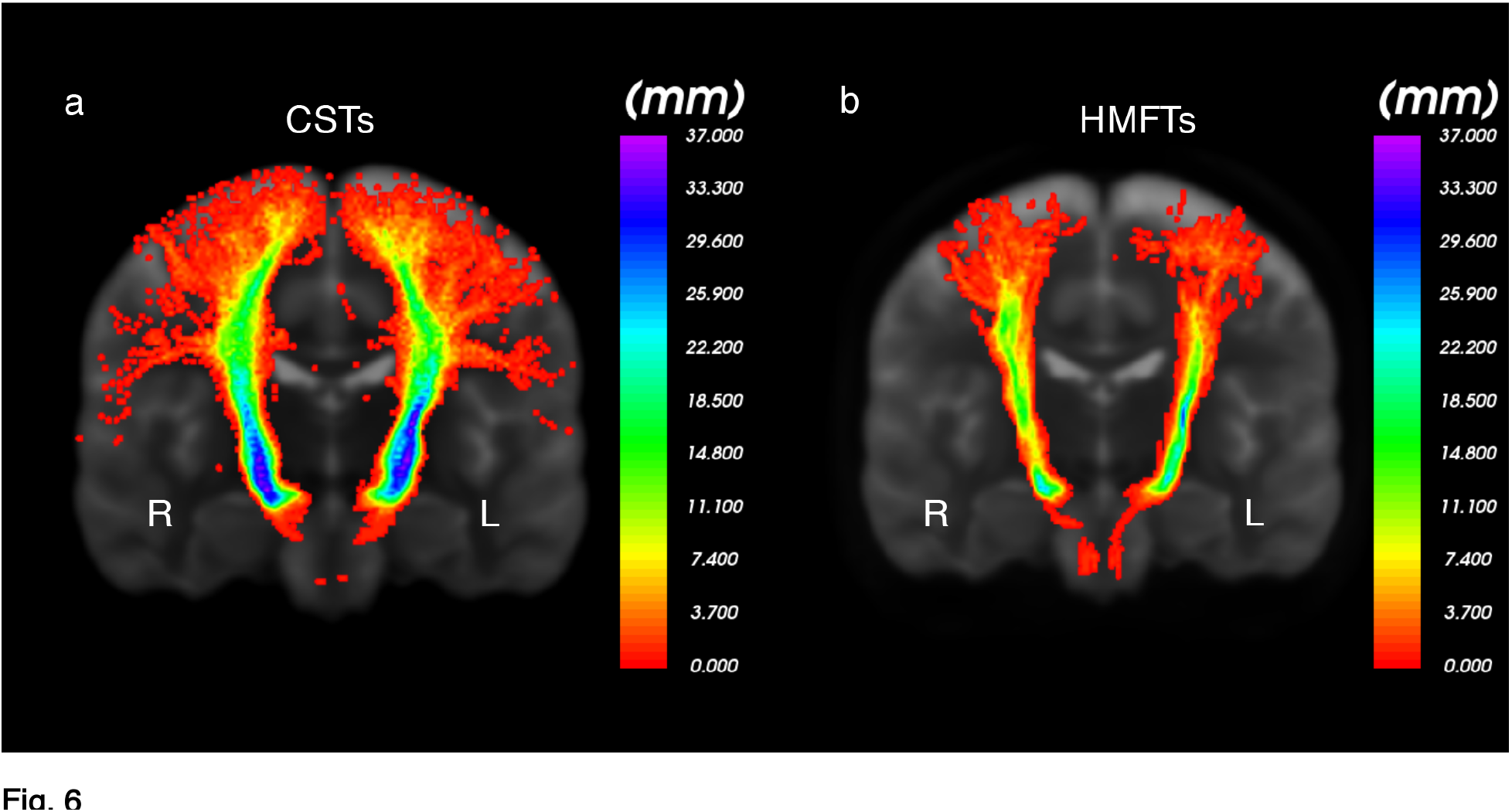
**a** Variability probabilistic heat map of the corticospinal tract (CST) in our 37 Washington University (WashU) healthy human subjects in the coronal plane. As indicated by the scale in the image, all or nearly all of our subjects had CST fibers passing from areas adjacent to the regions of interest (ROIs), with more variability in the cortical projections. Left is labeled as L and right as R. **b** Variability probabilistic heat map of hand-related motor fiber tracts (HMFTs) in our 37 WashU healthy human subjects in the coronal plane. Left is labeled as L and right as R. Despite the use of the same methodology on state-of-the-art data, the difference in variability is noticeable, potentially indicating limitations of the current tools in diffusion tensor imaging (DTI) image analysis. To the best of our knowledge, this is the first attempt at creating a variability probabilistic map of the hand motor fibers of healthy adults

### 3.5 Crossing fibers interference analysis

In an attempt to explain the apparent paucity of fibers in the hand motor area, we tested the hypothesis that crossing fibers at that anatomical level heavily affect the CST delineation, as described in the section 2.9. In agreement with our assumptions, the two sets of experiments described in section 2.9 indicated that there is an effect of the AF, SLFI, SLFII, SLFIII and CC4 (anterior midbody of the CC), as well as of the CC5 (posterior midbody of the CC), on the delineated HMFTs. Overall, based on the quantification analysis, there were no statistically significant correlations found between the volumes of the aforementioned tracts and the volumes of the HMFTs. We assume that this is because of the relatively small dataset used, as well as the limitations of our correlation method, which does not allow targeted quantification and spatial correlation of overlapping fibers only. On visual inspection, however, there is a clear spatial correspondence between those tracts in the overlapping areas. In particular, the presence of denser crossing fiber tracts in the area of the HMFTs, seemed to create “openings” within the HMFTs, possibly negatively affecting the density of the delineated HMFTs. The most relevant effect seemed to be due to the SLFII, which passes exactly through the hand motor area of interest, as well as the SLFIII. The AF seemed to interfere with the lateral part of the HMFTs, whereas the SLFI appeared to affect the medial part. The CC4 and CC5 heavily interfered with the anterior and posterior parts of the HMFTs, respectively. These effects are visually illustrated in the Fig. 7 and 8, where the interference of crossing fibers with the CST and, in particular with the HMFT, is clearly shown

**Fig. 7.**
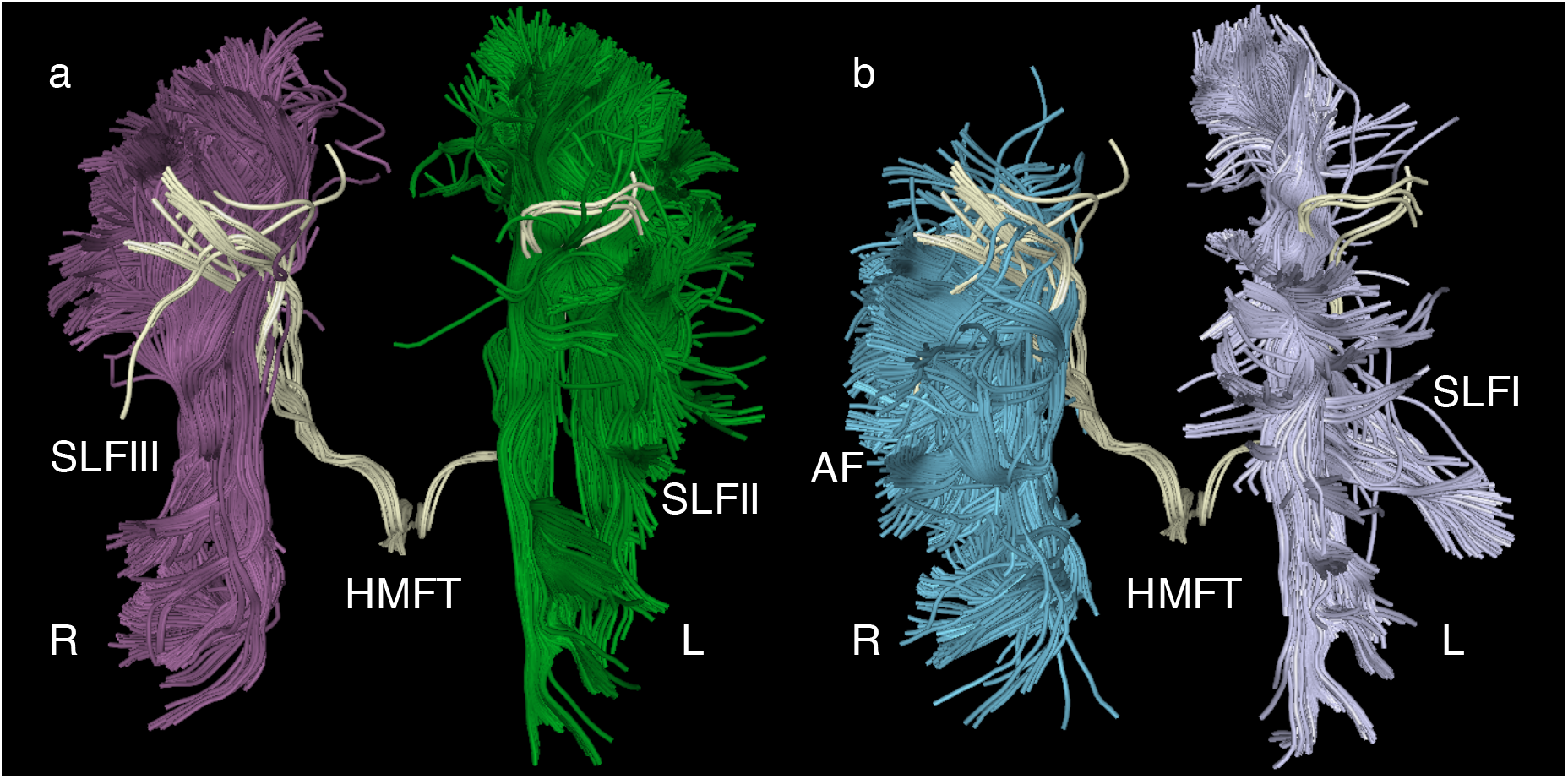
**a** Illustrative representation of the interference of superior longitudinal fascicle II (SLFII) (green) and SLFIII (purple) with the hand-related motor fiber tracts (HMFTs). This is an anterior view of an average case of a delineated HMFT (light yellow). SLFII and SLFIII appear to pass through the region of the corticospinal tract (CST) and in particular through the hand motor area. There are many “openings” within the HMFTs due to the passage of crossing fibers, which possibly affect the density of the delineated tract using diffusion magnetic resonance imaging (dMRI) tractography. **b** Depiction of the SLFI (gray) on the left and the arcuate fascicle (AF) (blue) on the right. AF seems to interfere with the HMFTs laterally, whereas SLFI appears to interfere medially. The right and left sides are marked as R and L, respectively, in all figures

**Fig. 8.**
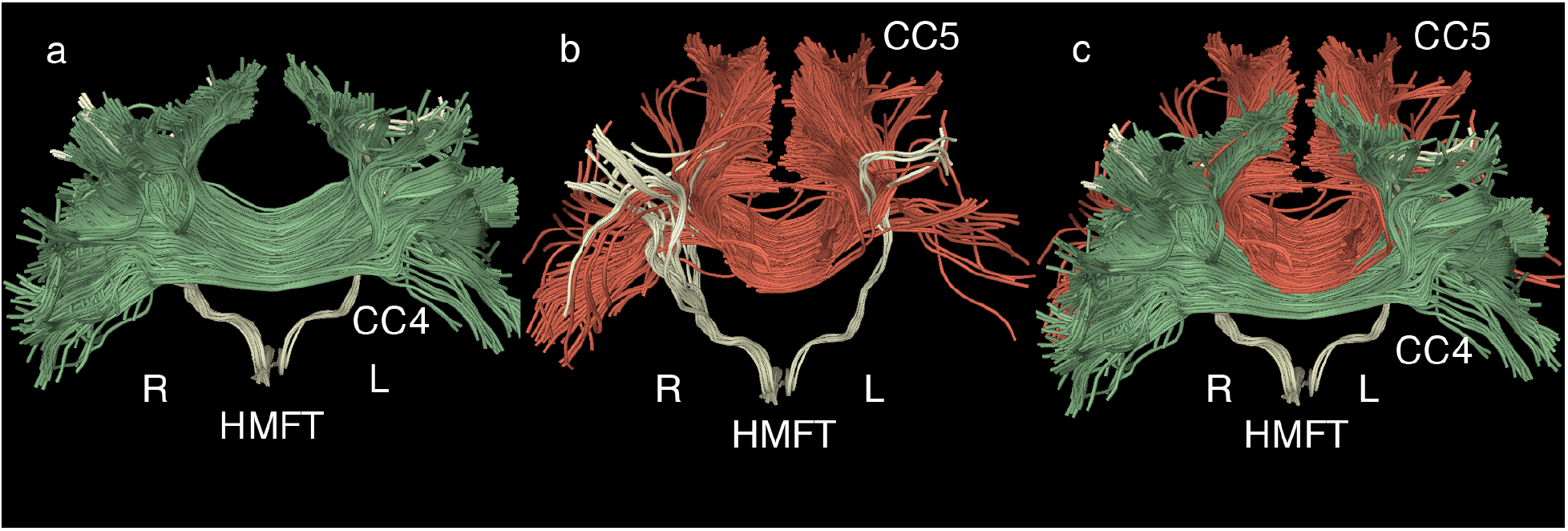
**a** Illustrative representation of the interference of the corpus callosum 4 (CC4, anterior midbody of the corpus callosum, in green) with the delineated hand-related motor fiber tracts (HMFTs). This is an anterior view of an average case of a delineated HMFT (light yellow). The entire HMFT seems to be affected by the crossing fibers, especially in its anterior aspect. **b** Depiction of the corpus callosum 5 (CC5, posterior midbody of the CC, in red) and its spatial correlation to the HMFTs (light yellow). This is the same anterior view of the HMFT. There is less interference than observed with respect to CC4, but the HMFTs’ density seems to be affected by the presence of crossing fibers in the posterior area. **c** Combined anterior representation of the previously illustrated CC4 (green) and CC5 (red) to show how heavily those fibers interfere with the HMFTs. All fibers are co-registered in the same Montreal Neurological Institute (MNI) 152 space. The right and left sides are marked as R and L, respectively, in all figures

## 4 Discussion

In the current study we delineated and quantified the corticospinal tract as a whole as well as its hand motor representation (HMFT) in 36 healthy human subjects derived from HCP DTI datasets. To this end, we applied two-tensor tractography, a higher-order dMRI-based tractographic procedure. Furthermore, we assessed inter-subject topographic variability of the CST and the HMFT in particular in this subject population. Inter-subject variability of the HMFTs was high as mapped using probabilistic heat maps, even though such variability was not as high for the CST.

The CST is probably the mostly quantitatively studied fiber tract in the human brain. Traditionally, it has been estimated as composed of approximately 1 million axons, which vary from 2-11 μm in diameter. Of these axons, approximately 700,000 are myelinated and 300,000 are unmyelinated (Ford and Hackney 1997; Lassek and Rasmussen 1939; Parent 1996; Yagishita et al. 1994). Several groups have employed combinatorial approaches using both direct brain electrostimulation and DTI tractography to delineate the CST (Berman et al. 2004, 2007; Kinoshita et al. 2005; Mikuni et al. 2007). A few groups have focused on using only DTI tractography in order to isolate the CST instead (Clark et al. 2003; Itoh et al. 2006; Mandelli et al. 2014; Okada et al. 2007; Petersen et al. 2016; Yamada et al. 2005). Based on their results, the more laterally located fibers corresponding to the motor area of the hand appeared to be poorly represented or not represented at all. Recent studies (Chen et al. 2016b; Farquharson et al. 2013; Mandelli et al. 2014) raised the issue of the poor sensitivity of the currently used DTI techniques in isolating the lateral part of the CST and suggested a potential advantage of using higher-order diffusion model tractography.

Our results are in agreement with other higher-order diffusion model tractography studies (Chen et al. 2016b; Mandelli et al. 2014; Mormina et al. 2015), suggesting that DTI tractography can be a robust tool for the successful delineation of the CST with many potential clinical applications. It should be noted that, to the best of our knowledge, currently there is no tractographic approach to delineate the CST in its entirety, given the interference of other prominent crossing fiber tracts. Nevertheless, active research from several groups is focused on the problem and the plan to reach a clinically translatable outcome (Pujol et al. 2015).

To the best of our knowledge, this is the first study to delineate exclusively the hand motor area of the CST and generate specific HMFTs variability maps in normal subjects. Our findings suggest that the fibers of SLF I, II and III and the AF interfere considerably with delineating the HMFTs, as shown in Fig. 4 and 7 (see e.g., https://www.ncbi.nlm.nih.gov/pmc/articles/PMC3163395/figure/F8/, https://www.ncbi.nlm.nih.gov/pmc/articles/PMC3163395/). Furthermore, crossing fibers of the CC body, CC subdivisions CC4 and CC5, seem to pose an additional obstacle to the precise evaluation of the descending HMFTs as shown in Fig. 8. The crossing fibers of these tracts appear to be the principal reason for the previously reported limitations of the tractographic algorithms to depict precisely the hand motor fibers of the CST. Supporting evidence for this view comes from studies on the clinical entity of congenital bilateral perisylvian syndrome (CBPS), a rare disease of congenital agenesis of the SLF (Bernal et al. 2010; Gropman et al. 1997; Kilinc et al. 2015; Lee et al. 2015; Saporta et al. 2011). DTI tractography performed in CBPS patients with SLF agenesis clearly isolates the hand area. Nevertheless, the trunk- and leg-related tracts could not be depicted, suggesting that other tracts might interfere with the delineation of these motor fibers (Kilinc et al. 2015). Based on our results as depicted in Fig. 6, we hypothesize that the occipitofrontal fascicle (OFF) interferes with the trunk- and leg-related fiber tracts (Nikos Makris et al. 2007). Further studies focused on those tracts are needed to confirm our hypothesis.

### 4.1 Clinical significance

In this study, we were able to identify and delineate the hand motor fiber tracts in 36 healthy subjects. Although the CST appeared to be reliably sampled, given its low spatial variation it seems there were notable limitations in delineating the hand motor area fiber representation. This is highly relevant for neurorehabilitation, neurology, child and developmental neurology, neurosurgery and neural engineering, and may provide valuable information to enable advancements in those fields.

#### 4.1.1 Clinical neurology/neurorehabilitation applications

Another major clinical gap that creates obstacles in the evaluation and follow-up of neurological patients is the fact that there is no available non-invasive technique that can be applied in everyday clinical practice to assess the integrity of nerve axons. Biophysical parameters such as FA value have been correlated with the integrity of nerves (Alshikho et al. 2016), and physical parameters such as number of fibers or tract volume have been correlated with nerve deficits associated with neurotrauma or disease (Feng et al. 2015). Several researchers have tried to use DTI tractography to assess the integrity of nerve fibers in several neurologic conditions such as stroke (Feng et al. 2015), ALS (Graaff et al. 2011), and MS (Filippi et al. 2016) and the results seem promising for clinical translation. Our methodology has demonstrated very good reproducibility in order to allow the assessment of the CST in different neurological conditions. The variability map of the CST and HMFTs that we created based on these healthy subjects can be extended further to provide a reliable and accurate reference for what is within normal range in the human population, allowing the early detection of pathologies. This provides a basis for screening neurological patients early in the course of a disorder associated with functional motor deterioration. Determining and quantifying the neural substrate of a motor deficit via our method will enable clinicians not only to monitor the progression of neurological conditions, but also to assess and quantify the effects of the therapeutic interventions and have a better understanding of the prognosis. Correlation of FA values with existing clinical scales can help develop new objective scales for assessing the integrity of tracts during the course of a disease and its treatment. Neurorehabilitation physicians, by contrast to neurosurgeons, can focus more on tract volume measurements in order to personalize physiotherapy schedules and timing and improve the accuracy of prognosis.

#### 4.1.2 Clinical translation of novel therapeutics in neuroscience

Neurologists, neurorehabilitation physicians and neural engineers are currently moving toward clinical translation of neural repair and neuroprotection strategies for CNS trauma and disease, such as TBI (Aertker et al. 2016), stroke (Manley et al. 2015), ALS (Petrou et al. 2016) and SCI (Lu et al. 2016). Nevertheless, in order to quantify and assess the effects of such novel therapeutics in neurological patients, non-invasive, reliable in vivo techniques are a key aspect of progressing to a safe clinical translation. Our method could easily be implemented in clinical studies in order to assess new therapeutic interventions that are progressing to clinical trials. This would allow objective quantification of the response of neural tissue to new treatments and in clinical trials.

## 5 Limitations and future studies

### 5.1 Study-specific limitations

Our dataset is a Human Connectome Project high-resolution dataset, meaning that our results may not be reproduced in lower resolution clinical datasets. Thus, further validation is needed with data sets of differing resolutions.

In addition, we chose to use a delineation methodology based on the drawing of manual ROIs in order to obtain the most accurate and clinically applicable results. For the purpose of the present study, this significantly increased accuracy and precision, which is important for establishing accurate variability maps in the healthy population. However, in future studies with large data cohorts (such as HCP with 1200 subjects) we may need to consider a machine-learning approach in order to facilitate the implementation of such methodology to make it feasible to perform such studies on a large scale. Although biophysical properties of diffusion imaging could be influenced by several factors such as age or sex, given that in this study we focused on topographical anatomical variability, we do not think our variability results would be affected by these factors. The choice of two-tensor UKF tractography might be highly useful for addressing some of the crossing-fiber tractography-related obstacles in order to allow better delineation of fiber tracts. However, the tractography is still limited to tracing only two-fibers and cannot trace fibers that pass through three-fiber crossings, which may contribute to increased variability of the tracts traced. Our future work will involve using more sophisticated algorithms that can address these shortcomings.

### 5.2 Technical considerations

Although we have selected datasets from human healthy subjects from the Connectome scanners considered to have the highest signal-to-noise ratio and resolution standards currently in the world, a future change in the resolution of acquisition or higher field strengths may affect the results (Alexander et al. 2001). Optimal parameters for visualizing tracts and nerve fibers remain to be developed.

There are well known and well described limitations to the tractography methods used in the present study. As mentioned in the discussion above, tract volume and FA values in DTI tractography can vary within and across subjects (Catani 2007; Heiervang et al. 2006), which necessitates that large numbers of subjects be studied to draw more accurate and clinically applicable conclusions. In addition, false positive or false negative results have been reported in tractography (Farquharson et al. 2013; Jones et al. 2013; O’Donnell and Pasternak 2015; O’Donnell and Westin 2011). Thus, we tried to ensure that our analysis was based on valid and reliable anatomic landmarks using manually drawn ROIs and a robust quality control analysis with optimization of the tractography algorithm parameters for the particular high-resolution dataset. Based on the significant problem of crossing fibers as discussed above, the two-tensor UKF tractography was chosen in order to minimize the effect of crossing fibers in our dataset as much as possible. Finally, a limitation of tractography especially relevant to neurosurgeons is that although it gives a good estimate of a fiber tract representation spatially, it provides neither the actual representation of the volume of the tract nor the number of fibers (Kinoshita et al. 2005; Nimsky et al. 2016). Thus, careful implementation and interpretation of tractography-based techniques is advised for neurosurgical applications.

## 6 Conclusions

We have demonstrated that two-tensor UKF tractography could be a robust tool for motor pathway isolation, in particular the CST and HFMT, significantly improving the tracking of tracts and overcoming certain intrinsic limitations of DTI tractography (i.e. crossing fibers or, as in other reports, peritumoral edema) as a higher-order model tractography algorithm. CST variability was delineated in healthy subjects using state-of-the-art Human Connectome Data in order to act as a reference for future anatomical studies and for establishing clinical correlations by neurorehabilitation physicians, neurologists, and neurosurgeons. The HMFTs were also isolated for the first time using solely DTI tractography. A healthy-subjects’ inter-subject variability heat map was described to allow future validation and improvement of the technique and imaging modalities and algorithms, given the great importance of this motor area for clinicians. Even though variance was much higher for the HMFTs than the CST, partially due to the presence of outliers, our methodology seems to be appropriate for the isolation of HMFTs in small group studies. Nevertheless, there remains significant need for improvement of the tools in order to ensure reliable single-subject analysis of the hand motor area in clinical settings. Further validation of the current results with a greater number of subjects is needed to establish an accurate variability map that can be used reliably for the clinical screening of fiber tract pathologies and to track neuroregenerative processes.

## Acknowledgements

We would like to thank the anonymous reviewers for providing useful comments on the manuscript. We would also like to thank Prof. Myron Spector for fruitful discussions and for his support.

## 7 Compliance with ethical standards

### 7.1 Funding

K.D. was partially supported by the Foundation for Education and European Culture (IPEP). M.T. was supported by the American Association of University Women (AAUW) and the Onassis Foundation. EY was supported by Colby College Research Fund 01 2836. NIH P41 EB015902. P.S. was supported by a NARSAD Young Investigator Award, grant number 22591 from the Brain and Behavior Research Foundation. N.M. was supported by R01AG042512 (National Institute of Aging & National Institute of Mental Health) and R21AT008865 (National Center for Complementary and Integrative Health).

### 7.3 Ethical approval

For this type of study formal consent is not required.

### 7.4 Informed consent

Informed consent was obtained from all individual participants included in the study.

## Abbreviations

CST: corticospinal tract
HMFTs: hand-related motor fiber tracts
PT: pyramidal tract
BA: Brodmann’s area
dMRI: diffusion magnetic resonance imaging
DTI: diffusion tensor imaging
HARDI: high angular resolution diffusion imaging
IC: internal capsule
SCI: spinal cord injury
TBI: traumatic brain injury
MS: Multiple sclerosis
ALS: Amyotrophic Lateral Sclerosis
AF: arcuate fascicle
SLF: superior longitudinal fascicle
CC: corpus callosum
HCP: Human Connectome Project
UKF deterministic tractography: unscented Kalman filter deterministic tractography
FA: fractional anisotropy
AD: axial diffusivity
RD: radial diffusivity
SD: standard deviation
WU-Minn HCP consortium: Washington University-University of Minnesota and Oxford University Human Connectome Project consortium
ROIs: regions of interest
WMQL: White Matter Query Language
AC: anterior commissure
SI: Symmetry index
WashU: Washington University
MNI: Montreal Neurological Institute
CBPS: congenital bilateral perisylvian syndrome
OFF: occipitofrontal fascicle

## References

Aertker, B. M., Bedi, S., & Cox Jr, C. S. (2016). Strategies for CNS repair following TBI. Experimental Neurology, 275, Part 3, 411–426. doi:10.1016/j.expneurol.2015.01.008

Alexander, A. L., Hasan, K. M., Lazar, M., Tsuruda, J. S., & Parker, D. L. (2001). Analysis of partial volume effects in diffusion-tensor MRI. Magnetic Resonance in Medicine, 45(5), 770–780.

Alshikho, M. J., Zürcher, N. R., Loggia, M. L., Cernasov, P., Chonde, D. B., Izquierdo Garcia, D., et al. (2016). Glial activation colocalizes with structural abnormalities in amyotrophic lateral sclerosis. Neurology, 87(24), 2554–2561. doi:10.1212/WNL.0000000000003427

Barnard, J. W., & Woolsey, C. N. (1956). A study of localization in the corticospinal tracts of monkey and rat. Journal of Comparative Neurology, 105(1), 25–50.

Basser, P. J. (2004). Scaling laws for myelinated axons derived from an electrotonic core-conductor model. Journal of Integrative Neuroscience, 3(2), 227–244.

Basser, P. J., Mattiello, J., & LeBihan, D. (1994). MR diffusion tensor spectroscopy and imaging. Biophysical journal, 66(1), 259.

Baumgartner, C. F., Michailovich, O., Levitt, J., Pasternak, O., Bouix, S., Westin, C.-F., & Rathi, Y. (2012). A unified tractography framework for comparing diffusion models on clinical scans. Presented at the CDMRI Workshop-MICCAI, Nice.

Berman, J. I., Berger, M. S., Chung, S., Nagarajan, S. S., & Henry, R. G. (2007). Accuracy of diffusion tensor magnetic resonance imaging tractography assessed using intraoperative subcortical stimulation mapping and magnetic source imaging. Journal of Neurosurgery, 107(3), 488–494. doi:10.3171/JNS-07/09/0488

Berman, J. I., Berger, M. S., Mukherjee, P., & Henry, R. G. (2004). Diffusion-tensor imaging—guided tracking of fibers of the pyramidal tract combined with intraoperative cortical stimulation mapping in patients with gliomas. Journal of Neurosurgery, 101(1), 66–72. doi:10.3171/jns.2004.101.1.0066

Bernal, B., Rey, G., Dunoyer, C., Shanbhag, H., & Altman, N. (2010). Agenesis of the arcuate fasciculi in congenital bilateral perisylvian syndrome: a diffusion tensor imaging and tractography study. Archives of Neurology, 67(4), 501–505. doi:10.1001/archneurol.2010.59

Bertrand, G., Blundell, J., & Musella, R. (1965). Electrical Exploration of the Internal Capsule and Neighbouring Structures During Stereotaxic Procedures. Journal of neurosurgery, 22(4), 333–343.

Betz, W. (1874). Anatomischer nachweis zweier gehirncentra. Zentralbl Med Wiss, 12(578,595).

Bouchard, C. (1866). Secondary degenerations of the spinal cord (1866). Translated into English by ER HUN (Utica, New York 1869). Cited by Lassek, Am: The pyramidal tract (Thomas, Springfield 1954).

Brothwell, D. R. (1960). THE ANTECEDENTS OF MAN. An Introduction to the Evolution of the Primates. By Le Gross Clark W. E.. Edinburgh University Press, 1959, pp. 374, 152 figures. 21s. Antiquity, 34(136), 307–307. doi:10.1017/S0003598X0003564X

Bucci, M., Mandelli, M. L., Berman, J. I., Amirbekian, B., Nguyen, C., Berger, M. S., & Henry, R. G. (2013). Quantifying diffusion MRI tractography of the corticospinal tract in brain tumors with deterministic and probabilistic methods. NeuroImage: Clinical, 3, 361–368. doi:10.1016/j.nicl.2013.08.008

Campbell, A. W. (1905). Histological Studies on the Localization of Cerebral Function. Cambridge Univ. Press. Cambridge. Mass.

Carlson, H. L., Laliberté, C., Brooks, B. L., Hodge, J., Kirton, A., Bello-Espinosa, L., et al. (2014). Reliability and variability of diffusion tensor imaging (DTI) tractography in pediatric epilepsy. Epilepsy & Behavior: E&B, 37, 116–122. doi:10.1016/j.yebeh.2014.06.020

Catani, M. (2007). From hodology to function. Brain, 130(3), 602–605. doi:10.1093/brain/awm008

Chen, Z., Tie, Y., Olubiyi, O., Zhang, F., Mehrtash, A., Rigolo, L., et al. (2016a). Corticospinal tract modeling for neurosurgical planning by tracking through regions of peritumoral edema and crossing fibers using two-tensor unscented Kalman filter tractography. International Journal of Computer Assisted Radiology and Surgery, 1–12. doi:10.1007/s11548-015-1344-5

Chen, Z., Tie, Y., Olubiyi, O., Zhang, F., Mehrtash, A., Rigolo, L., et al. (2016b). Corticospinal tract modeling for neurosurgical planning by tracking through regions of peritumoral edema and crossing fibers using two-tensor unscented Kalman filter tractography. International Journal of Computer Assisted Radiology and Surgery, 11(8), 1475–1486. doi:10.1007/s11548-015-1344-5

Chronister, R. B., & Hardy, S. G. P. (1997). The limbic system. Fundamental neuroscience. Churchill Livingstone, London, 443–454.

Clark, C. A., Barrick, T. R., Murphy, M. M., & Bell, B. A. (2003). White matter fiber tracking in patients with space-occupying lesions of the brain: a new technique for neurosurgical planning? NeuroImage, 20(3), 1601–1608. doi:10.1016/j.neuroimage.2003.07.022

Clarke, E., & O’Malley, C. D. (1996). The human brain and spinal cord: a historical study illustrated by writings from antiquity to the twentieth century. Norman Publishing. https://books.google.com/books?hl=en&lr=&id=Q_rO4ZFpUcgC&oi=fnd&pg=PR9&dq=the+human+brain+and+spinal+cord:+a+historical+study+illustrated+by+writtings+clarke+o%27malley&ots=rgeKMawCtC&sig=3G4xQ3JKY4t8NDJCCtEDLgulxVw. Accessed 22 May 2017

Cronbach, L. J. (1951). Coefficient alpha and the internal structure of tests. Psychometrika, 16(3), 297–334. doi:10.1007/BF02310555

Cruveilhier, J. (1853). Sur la paralysie musculaire progressive atrophique. Arch Gen Med, 91, 561–603.

Davidoff, R. A. (1990). The pyramidal tract. Neurology, 40(2), 332–332.

Dejerine, J. J., & Dejerine-Klumpke, A. (1895). Anatomie des centres nerveux (Vol. 1). Rueff. https://books-google-com.ezp-prod1.hul.harvard.edu/books?hl=en&lr=&id=xyMFrOTP9AYC&oi=fnd&pg=PA183&dq=Dejerine+1895&ots=jk7YjiVJmJ&sig=h47nB1pw1NrQ4LVENgBtAB54w3I. Accessed 31 August 2016

Dini, L., Vedolin, L., Bertholdo, D., Grando, R., Mazzola, A., Dini, S., et al. (2013). Reproducibility of quantitative fiber tracking measurements in diffusion tensor imaging of frontal lobe tracts: A protocol based on the fiber dissection technique. Surgical Neurology International, 4(1), 51. doi:10.4103/2152-7806.110508

Ellis, C. M., Suckling, J., Amaro Jr, E., Bullmore, E. T., Simmons, A., Williams, S. C. R., & Leigh, P. N. (2001). Volumetric analysis reveals corticospinal tract degeneration and extramotor involvement in ALS. Neurology, 57(9), 1571–1578.

Farquharson, S., Tournier, J.-D., Calamante, F., Fabinyi, G., Schneider-Kolsky, M., Jackson, G. D., & Connelly, A. (2013). White matter fiber tractography: why we need to move beyond DTI. Journal of Neurosurgery, 118(6), 1367–1377. doi:10.3171/2013.2.JNS121294

Fedorov, A., Beichel, R., Kalpathy-Cramer, J., Finet, J., Fillion-Robin, J.-C., Pujol, S., et al. (2012). 3D Slicer as an image computing platform for the Quantitative Imaging Network. Magnetic Resonance Imaging, 30(9), 1323–1341. doi:10.1016/j.mri.2012.05.001

Feng, W., Wang, J., Chhatbar, P. Y., Doughty, C., Landsittel, D., Lioutas, V.-A., et al. (2015). Corticospinal tract lesion load: An imaging biomarker for stroke motor outcomes. Annals of Neurology, 78(6), 860–870. doi:10.1002/ana.24510

Filippi, M., Pagani, E., Preziosa, P., & Rocca, M. A. (2016). The Role of DTI in Multiple Sclerosis and Other Demyelinating Conditions. In W. V. Hecke, L. Emsell, & S. Sunaert (Eds.), Diffusion Tensor Imaging (pp. 331–341). Springer New York. doi:10.1007/978-1-4939-3118-7_16

Fillard, P., Descoteaux, M., Goh, A., Gouttard, S., Jeurissen, B., Malcolm, J., et al. (2011). Quantitative evaluation of 10 tractography algorithms on a realistic diffusion MR phantom. NeuroImage, 56(1), 220–234. doi:10.1016/j.neuroimage.2011.01.032

Finger, S. (1994). Origins of neuroscience: a history of explorations into brain function. Oxford University Press.

Flechsig, P. E. (1877). Pyramidal tract in brain and cord. Archiv Heilkunde.

Flechsig, P. E. (1905). Einige Bemerkungen über die Untersuchungsmethoden der Grosshirnrinde, insbesondere des Menschen.

Ford, J. C., & Hackney, D. B. (1997). Numerical model for calculation of apparent diffusion coefficients (ADC) in permeable cylinders—comparison with measured ADC in spinal cord white matter. Magnetic Resonance in Medicine, 37(3), 387–394. doi:10.1002/mrm.1910370315

Galaburda, A. M., Corsiglia, J., Rosen, G. D., & Sherman, G. F. (1987). Planum temporale asymmetry, reappraisal since Geschwind and Levitsky. Neuropsychologia, 25(6), 853–868. doi:10.1016/0028-3932(87)90091-1

Gall, F. J., & Spurzheim, J. C. (1810). Anatomie et physiologie. Quoted in [7], 226.

Glasser, M. F., Sotiropoulos, S. N., Wilson, J. A., Coalson, T. S., Fischl, B., Andersson, J. L., et al. (2013). The minimal preprocessing pipelines for the Human Connectome Project. NeuroImage, 80, 105–124. doi:10.1016/j.neuroimage.2013.04.127

Graaff, M. M. van der, Sage, C. A., Caan, M. W. A., Akkerman, E. M., Lavini, C., Majoie, C. B., et al. (2011). Upper and extra-motoneuron involvement in early motoneuron disease: a diffusion tensor imaging study. Brain, 134(4), 1211–1228. doi:10.1093/brain/awr016

Gropman, A. L., Barkovich, A. J., Vezina, L. G., Conry, J. A., Dubovsky, E. C., & Packer, R. J. (1997). Pediatric congenital bilateral perisylvian syndrome: clinical and MRI features in 12 patients. Neuropediatrics, 28(4), 198–203. doi:10.1055/s-2007-973700

Heiervang, E., Behrens, T. E. J., Mackay, C. E., Robson, M. D., & Johansen-Berg, H. (2006). Between session reproducibility and between subject variability of diffusion MR and tractography measures. NeuroImage, 33(3), 867–877. doi:10.1016/j.neuroimage.2006.07.037

Hille, B. (2001). Ion Channels of Excitable Membranes (3rd edition.). Sinauer Associates Inc 2001-07.

Hirayama, K., Tsubaki, T., Toyokura, Y., & Okinaka, S. (1962). The representation of the pyramidal tract in the internal capsule and basis pedunculi. A study based on three cases of amyotrophic lateral sclerosis. Neurology, 12, 337.

Holmes, G., & May, W. P. (1909). On the exact origin of the pyramidal tracts in man and other mammals. SAGE Publications. http://journals.sagepub.com/doi/pdf/10.1177/003591570900200718. Accessed 23 May 2017

Holodny, A. I., Gor, D. M., Watts, R., Gutin, P. H., & Ulug, A. M. (2005). Diffusion-tensor MR tractography of somatotopic organization of corticospinal tracts in the internal capsule: Initial anatomic results in contradistinction to prior reports 1. Radiology, 234(3), 649–653.

Huang, H., Zhang, J., Jiang, H., Wakana, S., Poetscher, L., Miller, M. I., et al. (2005). DTI tractography based parcellation of white matter: Application to the mid-sagittal morphology of corpus callosum. NeuroImage, 26(1), 195–205. doi:10.1016/j.neuroimage.2005.01.019

Irfanoglu, M. O., Modi, P., Nayak, A., Hutchinson, E. B., Sarlls, J., & Pierpaoli, C. (2015). DR-BUDDI (Diffeomorphic Registration for Blip-Up blip-Down Diffusion Imaging) method for correcting echo planar imaging distortions. NeuroImage, 106, 284–299. doi:10.1016/j.neuroimage.2014.11.042

Itoh, D., Aoki, S., Maruyama, K., Masutani, Y., Mori, H., Masumoto, T., et al. (2006). Corticospinal Tracts by Diffusion Tensor Tractography in Patients With Arteriovenous Malformations. Journal of Computer Assisted Tomography, 30(4), 618–623.

Jones, D. K., Knösche, T. R., & Turner, R. (2013). White matter integrity, fiber count, and other fallacies: the do’s and don’ts of diffusion MRI. NeuroImage, 73, 239–254. doi:10.1016/j.neuroimage.2012.06.081

Kilinc, O., Ekinci, G., Demirkol, E., & Agan, K. (2015). Bilateral agenesis of arcuate fasciculus demonstrated by fiber tractography in congenital bilateral perisylvian syndrome. Brain & Development, 37(3), 352–355. doi:10.1016/j.braindev.2014.05.003

Kinoshita, M., Yamada, K., Hashimoto, N., Kato, A., Izumoto, S., Baba, T., et al. (2005). Fiber-tracking does not accurately estimate size of fiber bundle in pathological condition: initial neurosurgical experience using neuronavigation and subcortical white matter stimulation. NeuroImage, 25(2), 424–429. doi:10.1016/j.neuroimage.2004.07.076

Kuypers, H. (1958). Some projections from the peri-central cortex to the pons and lower brain stem in monkey and chimpanzee. Journal of Comparative Neurology, 110(2), 221–255.

Kuypers, H. G. (1964). The descending pathways to the spinal cord, their anatomy and function. Progress in brain research, 11, 178–202.

Lassek, A. M., & Rasmussen, G. L. (1939). The human pyramidal tract: a fiber and numerical analysis. Archives of Neurology & Psychiatry, 42(5), 872–876.

Le Bihan, D., Breton, E., Lallemand, D., Grenier, P., Cabanis, E., & Laval-Jeantet, M. (1986). MR imaging of intravoxel incoherent motions: application to diffusion and perfusion in neurologic disorders. Radiology, 161(2), 401–407.

Lee, D.-H., Park, J. W., Park, S.-H., & Hong, C. (2015). Have You Ever Seen the Impact of Crossing Fiber in DTI?: Demonstration of the Corticospinal Tract Pathway. PLoS ONE, 10(7). doi:10.1371/journal.pone.0112045

Levin, P. M., & Beadford, F. K. (1938). The exact origin of the cortico-spinal tract in the monkey. Journal of Comparative Neurology, 68(4), 411–422.

Lori, N. F., Akbudak, E., Shimony, J. S., Cull, T. S., Snyder, A. Z., Guillory, R. K., & Conturo, T. E. (2002). Diffusion tensor fiber tracking of human brain connectivity: aquisition methods, reliability analysis and biological results. NMR in Biomedicine, 15(7–8), 494–515.

Lu, P., Ahmad, R., & Tuszynski, M. H. (2016). Neural Stem Cells for Spinal Cord Injury. In M. H. Tuszynski (Ed.), Translational Neuroscience (pp. 297–315). Springer US. doi:10.1007/978-1-4899-7654-3_16

Mai, J. K., & Paxinos, G. (2011). The human nervous system. Academic Press. https://books.google.com/books?hl=en&lr=&id=J4cDpsl2rrQC&oi=fnd&pg=PP1&dq=The+Human+Nervous+System+3rd+Edition+mai+paxinos&ots=_VD68iKbhY&sig=UcvydF9sfxKiurf9NJ05eSeKzxM. Accessed 23 May 2017

Makris, N., Kennedy, D. N., McInerney, S., Sorensen, A. G., Wang, R., Caviness, V. S., & Pandya, D. N. (2005a). Segmentation of subcomponents within the superior longitudinal fascicle in humans: a quantitative, in vivo, DT-MRI study. Cerebral Cortex (New York, N.Y.: 1991), 15(6), 854–869. doi:10.1093/cercor/bhh186

Makris, N., Kennedy, D. N., McInerney, S., Sorensen, A. G., Wang, R., Caviness, V. S., & Pandya, D. N. (2005b). Segmentation of Subcomponents within the Superior Longitudinal Fascicle in Humans: A Quantitative, In Vivo, DT-MRI Study. Cerebral Cortex, 15(6), 854–869. doi:10.1093/cercor/bhh186

Makris, N., Meyer, J. W., Bates, J. F., Yeterian, E. H., Kennedy, D. N., & Caviness Jr., V. S. (1999). MRI-Based Topographic Parcellation of Human Cerebral White Matter and Nuclei: II. Rationale and Applications with Systematics of Cerebral Connectivity. NeuroImage, 9(1), 18–45. doi:10.1006/nimg.1998.0384

Makris, N., Papadimitriou, G. M., Sorg, S., Kennedy, D. N., Caviness, V. S., & Pandya, D. N. (2007). The occipitofrontal fascicle in humans: a quantitative, in vivo, DT-MRI study. NeuroImage, 37(4), 1100–1111. doi:10.1016/j.neuroimage.2007.05.042

Makris, N., Worth, A. J., Papadimitriou, G. M., Stakes, J. W., Caviness, V. S., Kennedy, D. N., et al. (1997). Morphometry of in vivo human white matter association pathways with diffusion-weighted magnetic resonance imaging. Annals of neurology, 42(6), 951–962.

Malcolm, J. G., Shenton, M. E., & Rathi, Y. (2010). Filtered multitensor tractography. IEEE transactions on medical imaging, 29(9), 1664–1675. doi:10.1109/TMI.2010.2048121

Mandelli, M. L., Berger, M. S., Bucci, M., Berman, J. I., Amirbekian, B., & Henry, R. G. (2014). Quantifying accuracy and precision of diffusion MR tractography of the corticospinal tract in brain tumors. Journal of Neurosurgery, 121(2), 349–358. doi:10.3171/2014.4.JNS131160

Manley, N. C., Azevedo-Pereira, R. L., Bliss, T. M., & Steinberg, G. K. (2015). Neural Stem Cells in Stroke: Intracerebral Approaches. In D. C. Hess (Ed.), Cell Therapy for Brain Injury (pp. 91–109). Springer International Publishing. doi:10.1007/978-3-319-15063-5_7

Mikuni, N., Okada, T., Enatsu, R., Miki, Y., Hanakawa, T., Urayama, S., et al. (2007). Clinical impact of integrated functional neuronavigation and subcortical electrical stimulation to preserve motor function during resection of brain tumors. Journal of Neurosurgery, 106(4), 593–598. doi:10.3171/jns.2007.106.4.593

Mori, S., Crain, B. J., Chacko, V. P., & Van Zijl, P. (1999). Three-dimensional tracking of axonal projections in the brain by magnetic resonance imaging. Annals of neurology, 45(2), 265–269.

Mormina, E., Longo, M., Arrigo, A., Alafaci, C., Tomasello, F., Calamuneri, A., et al. (2015). MRI Tractography of Corticospinal Tract and Arcuate Fasciculus in High-Grade Gliomas Performed by Constrained Spherical Deconvolution: Qualitative and Quantitative Analysis. American Journal of Neuroradiology, 36(10), 1853–1858. doi:10.3174/ajnr.A4368

Nimsky, C., Bauer, M., & Carl, B. (2016). Merits and Limits of Tractography Techniques for the Uninitiated. In J. Schramm (Ed.), Advances and Technical Standards in Neurosurgery (pp. 37–60). Springer International Publishing. doi:10.1007/978-3-319-21359-0_2

Norton, I., Essayed, W. I., Zhang, F., Pujol, S., Yarmarkovich, A., Golby, A., et al. (2017). SlicerDMRI: Open Source Diffusion MRI Software for Brain Cancer Research. Cancer Research.

Nyberg-Hansen, R., & Rinvik, E. (1963). Some comments on the pyramidal tract, with special reference to its individual variations in man. Acta Neurologica Scandinavica, 39(1), 1–30.

O’Donnell, L. J., & Pasternak, O. (2015). Does diffusion MRI tell us anything about the white matter? An overview of methods and pitfalls. Schizophrenia Research, 161(1), 133–141. doi:10.1016/j.schres.2014.09.007

O’Donnell, L. J., & Westin, C.-F. (2011). An introduction to diffusion tensor image analysis. Neurosurgery Clinics of North America, 22(2), 185–196, viii. doi:10.1016/j.nec.2010.12.004

Okada, T., Miki, Y., Kikuta, K., Mikuni, N., Urayama, S., Fushimi, Y., et al. (2007). Diffusion Tensor Fiber Tractography for Arteriovenous Malformations: Quantitative Analyses to Evaluate the Corticospinal Tract and Optic Radiation. American Journal of Neuroradiology, 28(6), 1107–1113. doi:10.3174/ajnr.A0493

Parent, A. (1996). Carpenter’s human neuroanatomy. Williams & Wilkins.

Peele, T. L. (1942). Cytoarchitecture of individual parietal areas in the monkey (Macaca mulatta) and the distribution of the efferent fibers. Journal of Comparative Neurology, 77(3), 693–737.

Pellegrino, R. G., Spencer, P. S., & Ritchie, J. M. (1984). Sodium channels in the axolemma of unmyelinated axons: a new estimate. Brain Research, 305(2), 357–360. doi:10.1016/0006-8993(84)90442-6

Petersen, M. V., Lund, T. E., Sunde, N., Frandsen, J., Rosendal, F., Juul, N., & Østergaard, K. (2016). Probabilistic versus deterministic tractography for delineation of the cortico-subthalamic hyperdirect pathway in patients with Parkinson disease selected for deep brain stimulation. Journal of Neurosurgery, 1–12. doi:10.3171/2016.4.JNS1624

Petrou, P., Gothelf, Y., Argov, Z., Gotkine, M., Levy, Y. S., Kassis, I., et al. (2016). Safety and Clinical Effects of Mesenchymal Stem Cells Secreting Neurotrophic Factor Transplantation in Patients With Amyotrophic Lateral Sclerosis: Results of Phase 1/2 and 2a Clinical Trials. JAMA neurology, 73(3), 337–344. doi:10.1001/jamaneurol.2015.4321

Pierpaoli, C., Barnett, A., Pajevic, S., Chen, R., Penix, L., Virta, A., & Basser, P. (2001). Water diffusion changes in Wallerian degeneration and their dependence on white matter architecture. Neuroimage, 13(6), 1174–1185.

Pitres, J. A. (1884). Recherches anatomo-cliniques sur les scléroses bilatérales de la moelle épinière consécutives à des lésions unilatérales du cerveau. G. Masson.

Pujol, S., Wells, W., Pierpaoli, C., Brun, C., Gee, J., Cheng, G., et al. (2015). The DTI Challenge: Toward Standardized Evaluation of Diffusion Tensor Imaging Tractography for Neurosurgery. Journal of Neuroimaging, 25(6), 875–882. doi:10.1111/jon.12283

Ravits, J., Appel, S., Baloh, R. H., Barohn, R., Brooks, B. R., Elman, L., et al. (2013). Deciphering amyotrophic lateral sclerosis: what phenotype, neuropathology and genetics are telling us about pathogenesis. Amyotrophic Lateral Sclerosis & Frontotemporal Degeneration, 14 Suppl 1, 5–18. doi:10.3109/21678421.2013.778548

Rieckmann, A., Van Dijk, K. R. A., Sperling, R. A., Johnson, K. A., Buckner, R. L., & Hedden, T. (2016). Accelerated decline in white matter integrity in clinically normal individuals at risk for Alzheimer’s disease. Neurobiology of Aging, 42, 177–188. doi:10.1016/j.neurobiolaging.2016.03.016

Ropper, A. H., Samuels, M. A., & Klein, J. (2014, May 25). Adams and Victor’s Principles of Neurology 10th Edition. McGraw-Hill Education. Accessed 19 August 2017

Russell, J. R., & DeMyer, W. (1961). The quantitative cortical origin of pyramidal axons of Macaca rhesus with some remarks on the slow rate of axolysis. Neurology, 11(2), 96–96.

Saporta, A. S. D., Kumar, A., Govindan, R. M., Sundaram, S. K., & Chugani, H. T. (2011). Arcuate fasciculus and speech in congenital bilateral perisylvian syndrome. Pediatric Neurology, 44(4), 270–274. doi:10.1016/j.pediatrneurol.2010.11.006

Schafer, E. A. (1883). Report on the lesions, primary and secondary, in the brain and spinal cord of the macacque monkey, exhibited by professors Ferrier and Yeo., iv, 316–26.

Schäfer, E. A. (1910). Experiments on the paths taken by volitional impulses passing from the cerebral cortex to the cord: The pyramids and the ventro-laterla descending tracts. Quarterly Journal of Experimental Physiology, 3(4), 355–373.

Schmahmann, J. D., & Pandya, D. N. (2006). Fiber Pathways of the Brain. OXFORD UNIVERSITY. Retrieved from https://www.researchgate.net/profile/Jeremy_Schmahmann/publication/230675466_Fiber_Pathways_of_the_Brain/links/00b7d5203ae2b4186e000000.pdf

Sherbondy, A. J., Dougherty, R. F., Napel, S., & Wandell, B. A. (2008). Identifying the human optic radiation using diffusion imaging and fiber tractography. Journal of Vision, 8(10), 12–12. doi:10.1167/8.10.12

Skirven, T. M., Osterman, A. L., Fedorczyk, J., & Amadio, P. C. (2011). Rehabilitation of the Hand and Upper Extremity, 2-Volume Set E-Book: Expert Consult. Elsevier Health Sciences.

Snow, N. J., Peters, S., Borich, M. R., Shirzad, N., Auriat, A. M., Hayward, K. S., & Boyd, L. A. (2016). A reliability assessment of constrained spherical deconvolution-based diffusion-weighted magnetic resonance imaging in individuals with chronic stroke. Journal of Neuroscience Methods, 257, 109–120. doi:10.1016/j.jneumeth.2015.09.025

Song, S.-K., Sun, S.-W., Ju, W.-K., Lin, S.-J., Cross, A. H., & Neufeld, A. H. (2003). Diffusion tensor imaging detects and differentiates axon and myelin degeneration in mouse optic nerve after retinal ischemia. NeuroImage, 20(3), 1714–1722.

Song, S.-K., Sun, S.-W., Ramsbottom, M. J., Chang, C., Russell, J., & Cross, A. H. (2002). Dysmyelination revealed through MRI as increased radial (but unchanged axial) diffusion of water. NeuroImage, 17(3), 1429–1436.

Stieltjes, B., Kaufmann, W. E., van Zijl, P. C., Fredericksen, K., Pearlson, G. D., Solaiyappan, M., & Mori, S. (2001). Diffusion tensor imaging and axonal tracking in the human brainstem. NeuroImage, 14(3), 723–735. doi:10.1006/nimg.2001.0861

Tuch, D. S., Reese, T. G., Wiegell, M. R., Makris, N., Belliveau, J. W., & Wedeen, V. J. (2002). High angular resolution diffusion imaging reveals intravoxel white matter fiber heterogeneity. Magnetic Resonance in Medicine, 48(4), 577–582.

Tukey, J. W. (1977). Exploratory Data Analysis. (1 edition.). Reading, Mass: Pearson.

Turck, L. (1851). Über den Zustand der Sensibilität nach teilweiser Trennung des Rückenmarks. Zeit. Der k. R. Gesellschaft der Aerzte zu Wien, 189.

Uğurbil, K., Xu, J., Auerbach, E. J., Moeller, S., Vu, A., Duarte-Carvajalino, J. M., et al. (2013). Pushing spatial and temporal resolution for functional and diffusion MRI in the Human Connectome Project. NeuroImage, 80, 80–104. doi:10.1016/j.neuroimage.2013.05.012

Walker, L., Chang, L.-C., Nayak, A., Irfanoglu, M. O., Botteron, K. N., McCracken, J., et al. (2016). The diffusion tensor imaging (DTI) component of the NIH MRI study of normal brain development (PedsDTI). NeuroImage, 124, Part B, 1125–1130. doi:10.1016/j.neuroimage.2015.05.083

Wassermann, D., Makris, N., Rathi, Y., Shenton, M., Kikinis, R., Kubicki, M., & Westin, C.-F. (2013). On Describing Human White Matter Anatomy: The White Matter Query Language. Medical image computing and computer-assisted intervention: MICCAI … International Conference on Medical Image Computing and Computer-Assisted Intervention, 16(0 1), 647–654.

Wassermann, D., Makris, N., Rathi, Y., Shenton, M., Kikinis, R., Kubicki, M., & Westin, C.-F. (2016). The white matter query language: a novel approach for describing human white matter anatomy. Brain Structure & Function. doi:10.1007/s00429-015-1179-4

Willis, T. (1965). Cerebri Anatome, London 1664, englished 1681 by Samuel Pordage. Idem, The anatomy of the brain an nerves, 2.

Yagishita, A., Nakano, I., Oda, M., & Hirano, A. (1994). Location of the corticospinal tract in the internal capsule at MR imaging. Radiology, 191(2), 455–460.

Yamada, K., Kizu, O., Ito, H., Kubota, T., Akada, W., Goto, M., et al. (2005). Tractography for arteriovenous malformations near the sensorimotor cortices. AJNR. American journal of neuroradiology, 26(3), 598–602.

Yousry, T. A., Schmid, U. D., Alkadhi, H., Schmidt, D., Peraud, A., Buettner, A., & Winkler, P. (1997). Localization of the motor hand area to a knob on the precentral gyrus. A new landmark. Brain, 120(1), 141–157. doi:10.1093/brain/120.1.141

